# A metabolically controlled contact site between lipid droplets and vacuoles

**DOI:** 10.1101/2023.04.21.537652

**Authors:** Duy Trong Vien Diep, Javier Fernández Collado, Marie Hugenroth, Mike Wälte, Oliver Schmidt, Ruben Fernández-Busnadiego, Maria Bohnert

**Affiliations:** Institute of Cell Dynamics and Imaging, University of Münster, Von-Esmarch-Str. 56, 48149 Münster, Germany; Cells in Motion Interfaculty Centre (CiM), University of Münster, Germany; Institute of Neuropathology, University Medical Center Göttingen, 37099, Göttingen, Germany; Institute of Cell Biology, Biocenter Innsbruck, Medical University of Innsbruck, 6020 Innsbruck, Austria; Cluster of Excellence “Multiscale Imaging: from Molecular Machines to Networks of Excitable Cells” (MBExC), University of Göttingen, Göttingen, Germany

**Keywords:** Ldo16, Ldo45, LDAF1, promethin, Vac8, Pdr16, seipin, Sei1, Ldb16, Lipophagy

## Abstract

The lipid <u>d</u>roplet (LD) <u>o</u>rganization proteins Ldo16 and Ldo45 affect multiple aspects of LD biology in yeast. They are linked to the LD biogenesis machinery seipin, and their loss causes defects in LD positioning, protein targeting, and breakdown. However, their molecular roles remained enigmatic. Here we report that Ldo16/45 form a tether-complex with Vac8 for creation of <u>v</u>a<u>c</u>uole <u>lip</u>id droplet (vCLIP) contact sites, which can form in the absence of seipin. The phosphatidylinositol transfer protein Pdr16 is a further vCLIP-resident recruited by Ldo45. While only an LD-subpopulation is engaged in vCLIPs at glucose-replete conditions, nutrient stress results in vCLIP expansion, and vCLIP defects impair lipophagy upon prolonged starvation. In summary, Ldo16/45 are multifunctional proteins that orchestrate formation of a metabolically-regulated contact site. Our studies suggest an unexpected link between LD biogenesis and breakdown, and open the door to a deeper understanding of how lipid homeostasis is maintained during metabolic challenges.

## INTRODUCTION

Lipids function as major components of biological membranes, as highly efficient energy reservoir, and as signaling molecules. Lipid droplets (LDs) are ubiquitous organelles that are specialized in lipid storage in the form of neutral lipids, and actively involved in diverse lipid metabolic pathways. Having a dedicated lipid storage system available is important, both to maintain a continuous supply of lipids when exogenous nutrients are scarce, and to prevent the accumulation of toxic lipid species. Through their ability to safely store variable amounts of neutral lipids, LDs thus effectively act as lipid buffers that enable the cell to handle fluctuations in lipid availability, and to maintain lipid homeostasis.

LDs undergo cycles of biogenesis and breakdown dependent on the metabolic state of the cell (Hashemi and Goodman, 2015; Henne et al., 2018). Their formation starts at the endoplasmic reticulum (ER) with the synthesis of esterified neutral lipids, such as triacylglycerol and sterol esters (Farese and Walther, 2022). Newly formed neutral lipids are released into the phospholipid bilayer bordering the ER, where they are soluble at low concentrations, but form lipid lenses at higher concentrations (Thiam and Ikonen, 2021). These lenses grow and ultimately bud toward the cytosol, resulting in mature LDs that consist of a neutral lipid core compartment covered by a phospholipid monolayer derived from the cytosolic leaflet of the ER membrane (Olzmann and Carvalho, 2019). The LD surface is decorated with a dedicated set of proteins. These include a number of lipid metabolic enzymes involved in neutral lipid synthesis and breakdown, and in the synthesis of membrane lipids (Olarte et al., 2022). LD number and/or size increase at conditions where excess lipids accumulate inside the cell. In contrast, when membranes need to quickly expand, or when large amounts of lipids have to be degraded for ATP generation, lipid molecules are liberated from LDs. Liberation of lipids occurs by two alternative routes, lipolysis and lipophagy. During lipolysis, cytosolic lipases bind to LDs and hydrolyze triacylglycerol and sterols esters into their building blocks (Athenstaedt and Daum, 2003, 2005; Athenstaedt et al., 1999; Kurat et al., 2006). Lipophagy on the other hand is an organelle-specific type of autophagy where LDs are internalized into the lysosome and degraded by lysosomal lipases (Seo et al., 2017; Tsuji et al., 2017; Vevea et al., 2015; Wang et al., 2014a; van Zutphen et al., 2014). To maintain lipid homeostasis particularly during times of changing nutrient availability, the processes of LD formation and breakdown need to be tightly controlled, but the underlying molecular mechanisms are not fully understood.

The lipid <u>d</u>roplet <u>o</u>rganization proteins of 16 and 45 kDa (Ldo16 and Ldo45) have been recently identified as novel LD surface proteins in *Saccharomyces cerevisiae* (from here on yeast), which affect multiple aspects of the LD life cycle (Eisenberg-Bord et al., 2018; Teixeira et al., 2018). These proteins are derived from overlapping genes, so that the smaller LDO variant Ldo16 is identical to the C-terminus of the longer Ldo45 (Figure 1A, bottom; Figure S1A) (Bohnert, 2020). LDAF1/promethin in humans and dmLDAF1 in fly are proteins with structural similarities to LDO, which also localize to LDs (Castro et al., 2019; Chartschenko et al., 2021; Chung et al., 2019). The LDO proteins as well as their putative metazoan counterparts LDAF1/promethin and dmLDAF1 are physically and functionally linked to seipin (Castro et al., 2019; Chartschenko et al., 2021; Chung et al., 2019), an evolutionarily conserved LD biogenesis machinery (Arlt et al., 2022; Fei et al., 2008; Grippa et al., 2015; Kim et al., 2022; Klug et al., 2021; Prasanna et al., 2021; Renne et al., 2022; Salo et al., 2016, 2019; Sui et al., 2018; Szymanski et al., 2007; Wang et al., 2016; Yan et al., 2018; Zoni et al., 2021). Mutations in seipin result in Berardinelli Seip congenital generalized lipodystrophy type 2, and in neurological defects (Magré et al., 2001; Windpassinger et al., 2004). The yeast LDO proteins are not uniformly distributed across the cellular LD pool, but instead enriched on an LD subpopulation located adjacent to the nucleus vacuole junction (NVJ), a contact site between the nuclear ER and the vacuole (yeast lysosome) (Eisenberg-Bord et al., 2018). Ldo45, but not Ldo16, is required for association of the phosphatidylinositol transfer protein (PITP) Pdr16 to the surface of these LDs (Eisenberg-Bord et al., 2018; Teixeira et al., 2018). Similarly, fly dmLDAF1 localizes to an LD subpopulation (Chartschenko et al., 2021). The LDO proteins are also involved in nutrient stress responses. While LDs in logarithmically growing cells cultured in glucose-rich medium are dispersed through the cell, they accumulate at the NVJ during the diauxic shift, when cells run out of glucose (Barbosa et al., 2015; Eisenberg-Bord et al., 2018; Hariri et al., 2018), and this response is blocked in the absence of LDO proteins (Eisenberg-Bord et al., 2018). If cells are further cultured into late stationary phase, LDs are degraded by lipophagy, a process that is prevented in *ldo* mutants (Teixeira et al., 2018). Despite these diverse LDO-related phenotypes, the molecular functions of the LDO proteins remained unclear.

**Figure 1:**
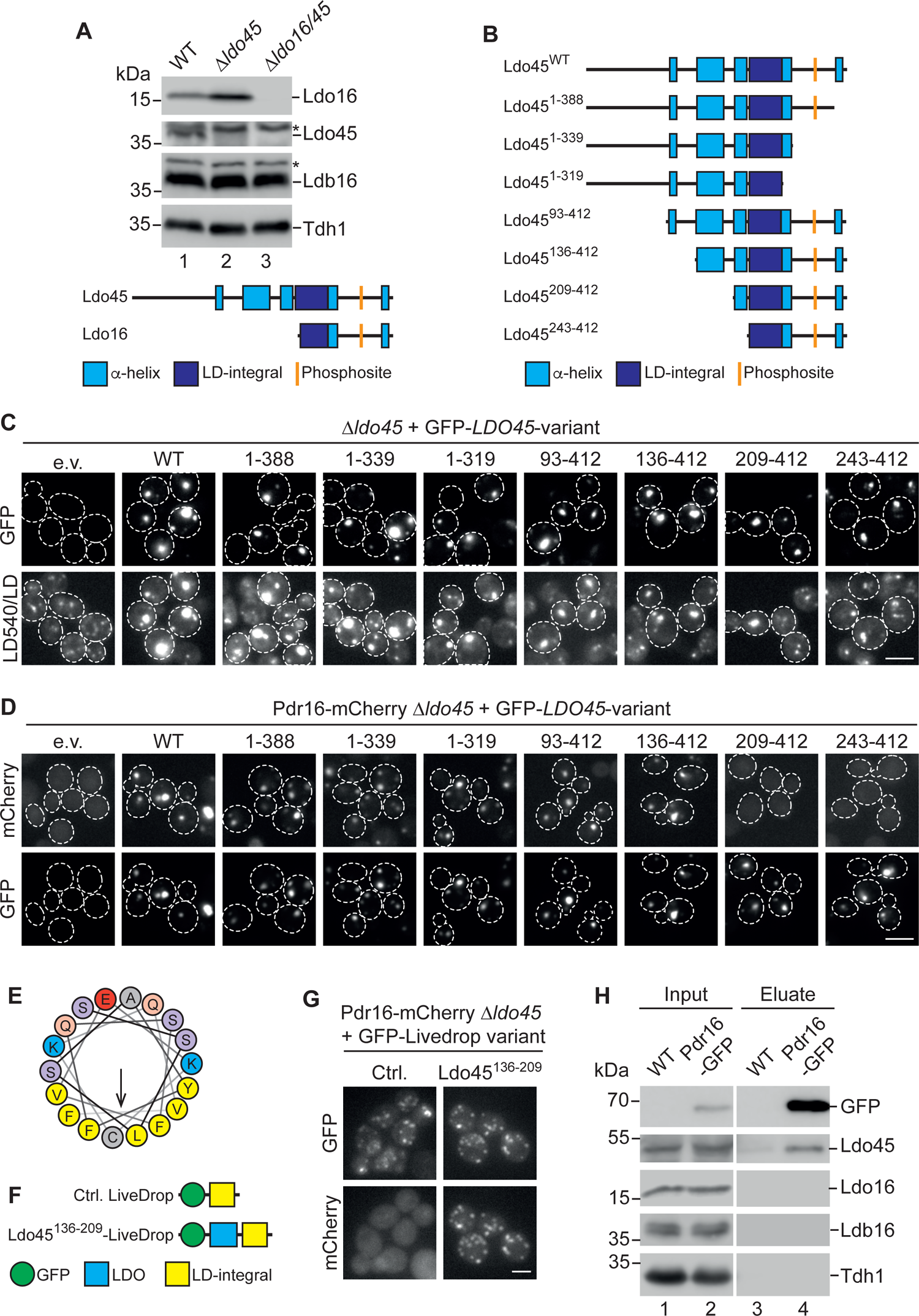
An amphipathic helix in the Ldo45 N-terminal cytosolic part mediates Pdr16 targeting to LDs. (A) Top: Indicated strains were cultured to logarithmic growth phase in synthetic medium containing 2% glucose, subjected to protein extraction, and analyzed by SDS-PAGE and Western blotting. Asterisks indicate unspecific bands. Bottom: Schematic representation of Ldo45 and Ldo16. Predicted α-helical regions in light blue, putative LD-integral hydrophobic α-helical domains in dark blue, putative phosphosites at position S366 (Ldo45) and S102 (Ldo16) in orange. (B) Schematic representation of native Ldo45 (Ldo45^WT^) and of truncated Ldo45 variants. Numbers indicate amino acids present in Ldo45 variants. Secondary structure elements were deleted as indicated. (C) Indicated GFP-Ldo45 variants were expressed under control of a *TEF2* promoter from centromeric plasmids in cells with a genomic *LDO45* deletion. Cells were cultured to logarithmic growth phase in synthetic medium containing 2% glucose, stained with the neutral lipid dye LD540, and analyzed by fluorescence microscopy. e.v., empty vector. Scale bar, 5 µm. (D) 1′*ldo45* Pdr16-mCherry were analyzed as described in (C), with the exception that LD540 staining was omitted. Scale bar, 5 µm. (E) Helical wheel projection of Ldo45 residues 137-154 shows amphipathic character of this α-helical stretch. Hydrophobic residues in yellow. (F) Schematic representation of standard LiveDrop (top) and Ldo45^136-209^-LiveDrop (bottom), a variant carrying an amphipathic region of Ldo45. (G) Pdr16-mCherry 1′*ldo45* cells expressing LiveDrop variants described in (F) from a centromeric plasmid under control of the *TEF2* promoter were cultured to logarithmic growth phase in synthetic medium containing 2% glucose and analyzed by fluorescence microscopy. Ctrl., control. Scale bar, 5 µm. (H) Pdr16-GFP and untagged control cells were cultured to logarithmic growth phase in synthetic medium containing 2% glucose and subjected to immunoprecipitation using GFP-trap agarose, followed by SDS-PAGE and Western blotting. Input, 5%, Eluate, 100%.

Here, we report that in addition to their established function as partner proteins of the seipin LD biogenesis machinery, the LDO proteins form extensive <u>v</u>a<u>c</u>uole <u>lip</u>id droplet (vCLIP) contact sites (also see Álvarez-Guerra et al., submitted in parallel to this study). vCLIP formation does not depend on the presence of functional seipin. LD-vacuole tethering is mediated by the C-terminal cytosolic domain that is present in both Ldo16 and Ldo45, and by the vacuolar adaptor protein Vac8. The Ldo45-specific N-terminal cytosolic domain in turn recruits the PITP Pdr16 to vCLIP. LD-vacuole contact sites are metabolically regulated, and are a prerequisite for the process of LD breakdown by lipophagy. We conclude that Ldo16/45 are multifunctional proteins with central roles in lipid homeostasis.

## RESULTS

### LD targeting of the phosphatidylinositol transfer protein Pdr16 is mediated by an amphipathic helix in the Ldo45 N-terminal cytosolic domain

So far, the functions of the LDO proteins Ldo16 and Ldo45 have been studied in deletion mutants lacking either one or both proteins (Eisenberg-Bord et al., 2018; Teixeira et al., 2018). The proteins are encoded by overlapping genomic regions, and hence deletion of a fragment corresponding to the *LDO16* coding region results in loss of both Ldo16 and Ldo45. We therefore refer to this mutant as Δ*ldo16/45* (Figure 1A, lane 3; Figure S1A and B). Deletion of a fragment in the 5’ region of the *LDO45* coding region on the other hand blocks formation of Ldo45 (Figure 1A, lane 2; Figure S1A), while Ldo16 is efficiently synthesized, resulting in elevated Ldo16 levels when compared to controls (Figure 1A, lanes 1 and 2), and we term this mutant Δ*ldo45.* To obtain a more detailed view on the roles of the two LDO variants, we generated a set of plasmids for expression of truncated LDO proteins designed to map the subdomains responsible for the different LDO-related LD phenotypes. Ldo45 contains a central hydrophobic α-helical stretch that is likely integrated into the LD phospholipid monolayer, and exposes hydrophilic N- and C-terminal domains to the cytosol (Eisenberg-Bord et al., 2018; Teixeira et al., 2018). The shorter LDO variant Ldo16 corresponds to the C-terminal region of Ldo45, comprising a part of the hydrophobic LD-integral domain, and the cytosolic C-terminal domain (Figure 1A, bottom). We generated three Ldo45 variants with truncated C-termini: Ldo45^1-388^ lacks the C-terminal 24 amino acids that according to secondary structure predictions could be α-helical; Ldo45^1-339^ additionally lacks almost the complete, largely disordered cytosolic C-terminal domain, except for the 20 amino acids adjacent to the LD-integral domain, which are predicted to form an α-helical extension of the putative LD-integral domain; and Ldo45^1-319^ lacks the entire cytosolic C-terminus. As for N-terminally truncated variants, we generated Ldo45^93-412^, which lacks the very N-terminal, putatively disordered part, and Ldo45^136-412^, Ldo45^209-412^, and Ldo45^243-412^, in which three cytosolic α-helixes were additionally successively deleted (Figure 1B). All Ldo45 variants were fused to an N-terminal GFP tag for detection, and expressed from a strong *TEF2* promoter. When transformed into *ldo45*1′ cells, all truncation variants localized to LDs, labeled by the neutral lipid dye LD540, similar to the wild-type protein (Figure 1C). Of note, all variants induced, to a variable degree, formation of LD clusters, similar to what has been reported before for overexpression of full length Ldo45 (Figure 1C, bottom row, compare to empty vector control) (Eisenberg-Bord et al., 2018; Teixeira et al., 2018).

Ldo45 was previously shown to be required for targeting of the PITP Pdr16 to LDs (Eisenberg-Bord et al., 2018; Teixeira et al., 2018). To map the region of Ldo45 responsible for this function, all Ldo45 truncation variants were expressed in *ldo45*1′ Pdr16-mCherry cells. While all C-terminally truncated Ldo45 variants efficiently supported Pdr16 targeting to LDs, the N-terminal truncations showed differential effects. Variants Ldo45^93-412^ and Ldo45^136-412^ mediated Pdr16 targeting similar to full length Ldo45. In contrast, Ldo45^209-412^ and Ldo45^243-412^ completely lost this function, resulting in a dispersed Pdr16 localization in the cytosol comparable to the phenotype observed in cells lacking *LDO45* completely (Figure 1D). Similar results were obtained with plasmids expressing untagged Ldo45 variants from the endogenous *LDO45* promoter (Figure S1C). This indicated that the central α-helix in the Ldo45 N-terminus, corresponding to amino acids 136-209, might be important for Pdr16 localization. Structural predictions revealed that large parts of this domain have an amphipathic character (exemplified in Figure 1E for amino acids 137-154). To test whether this domain is sufficient for Pdr16 targeting, we generated a synthetic construct consisting of the LD probe LiveDrop (Wang et al., 2016), and amino acids 136-209 of Ldo45 (Figure 1F). This construct, GFP-Ldo45^136-209^-LiveDrop, was targeted to LDs like regular LiveDrop (Figure S1D). When expressed in cells lacking *LDO45*, GFP-Ldo45^136-209^-LiveDrop mediated Pdr16 binding to LDs, while regular GFP-LiveDrop had no effect (Figure 1G). This finding suggested that Ldo45 might directly interact with Pdr16 and recruit the PITP to the LD surface. We therefore performed GFP-trap immunoprecipitation with cells expressing Pdr16-GFP. Using an antibody directed against native LDO proteins, we found that specifically Ldo45 was co-isolated with Pdr16-GFP, while Ldo16 as well as the seipin subunit Ldb16 were not co-isolated (Figure 1H, lane 4). In summary, an amphipathic α-helix in the N-terminal soluble domain of Ldo45 is responsible for Pdr16 targeting to LDs. This is consisten with the previous finding that Ldo16 cannot functionally replace Ldo45, as this domain is not present in the Ldo16 protein.

### The C-terminal cytosolic domain of Ldo16/45 has a role in nutrient stress-induced LD accumulation at the NVJ

We previously found that *ldo* mutants simultaneously lacking both Ldo16 and 45 fail to accumulate LDs at the NVJ during the diauxic shift, when cells run out of glucose (Eisenberg-Bord et al., 2018). We wondered which aspect of the physiological changes that occur during the diauxic shift is responsible for this phenotype, and subjected cells to a glucose restriction protocol, where we cultured them on medium supplemented with 2% glucose overnight, shifted them to medium containing only 0.001% glucose, and cultured them for a further 4 hours. This treatment efficiently induced LD accumulation at the NVJ in control cells, while LDs in *ldo16/45*1′ cells encircled the nucleus (Figure 2A and Figure S2A). Glucose restriction was therefore used as a simple and robust protocol to characterize the role of the LDO proteins in LD accumulation at the NVJ in more detail. Using plasmids for expression of either Ldo16 or Ldo45 alone from their endogenous promoters, we next tested which of the LDO proteins was responsible for this effect, and found that both proteins individually efficiently supported LD accumulation at the NVJ (Figure 2B and Figure S2B). To map the part of the protein required for this effect, we focused on the shorter, structurally simpler Ldo16. We generated plasmids for expression of three Ldo16 truncation variants lacking parts of the C-terminal cytosolic domain, Ldo16^1-124^, Ldo16^1-75^ and Ldo16^1-55^ (Figure 2C). GFP-tagged Ldo16^1-124^ and Ldo16^1-75^ colocalized with the neutral lipid dye LD540 similar to full length Ldo16, demonstrating targeting to LDs (Figure S2C). In contrast, Ldo16^1-55^ localized to foci distinct from LDs (Figure S2C), suggesting that amino acids 56-75, which form another amphipathic helix, are important for Ldo16 targeting. Due to its mislocalization, Ldo16^1-55^ was excluded from further investigations. When subjecting cells expressing the remaining Ldo16 variants to glucose restriction, we found that while Ldo16^1-124^ partially supported LD accumulation at the NVJ, Ldo16^1-75^ was virtually inactive, resulting in a phenotype close to that of Δ*ldo16/45* cells (Figure 2D and Figure S2D). Of note, phosphoproteomic studies have in the past detected a phosphosite at serine 102 of Ldo16, and hence within the determinant for NVJ accumulation of LDs (Albuquerque et al., 2008; Lanz et al., 2021; Pultz et al., 2012; Swaney et al., 2013; Zhou et al., 2021). We therefore performed Phos-tag SDS-PAGE of cells expressing wild-type Ldo16 using an antibody directed against Ldo16, and indeed detected a mobility shifted protein fraction, indicating a potential phosphorylation (Figure 2E, lane 2). We then generated a plasmid pLdo16^S102A^, where the putatively phosphorylated serine was replaced by an alanine. Ldo16^S102A^ exclusively migrated at the position of unphosphorylated Ldo16, showing that the shifted fraction indeed reflected phosphorylation at position 102 (Figure 2E, lane 3). As our Ldo16 antibody showed several cross-reactive bands, we analyzed cells expressing Ldo16/45 fused to a C-terminal FLAG tag, which also resolved a mobility-shifted Ldo16 species. Additionally, we detected one prominent and several weaker bands migrating above Ldo45, suggesting that the longer LDO protein might be phosphorylated as well (Figure 2E, lane 5). We next generated two phosphomimetic variants Ldo16^S102E^ and Ldo16^S102D^, and tested their ability to support LD accumulation at the NVJ, alongside the non-phosphorylatable mutant Ldo16^S102A^. We found that upon expression of the non-phosphorylatable and the phosphomimetic variants, LDs efficiently accumulated at the NVJ in response to glucose restriction (Figure 2F and Figure S2E), suggesting that the phosphorylation state of Ldo16 is not critical for this effect. As the LDO proteins are seipin partner proteins (Eisenberg-Bord et al., 2018; Teixeira et al., 2018), we next individually deleted *SEI1* and *LDB16*, to test whether seipin was also required for LD accumulation at the NVJ. While loss of seipin resulted in LD clustering as described before (Fei et al., 2008; Szymanski et al., 2007), these LD clusters still accumulated adjacent to vacuolar membranes (Figure 2G), suggesting a seipin-independent function of the LDO proteins. Finally, to obtain high resolution information on the LDs positioned adjacent to the NVJ, we analyzed glucose restricted cells by cryo-electron tomography of vitrified wild-type cells (Collado et al., 2019). We detected numerous LDs adjacent to nuceus-vacuole interfaces in glucose restricted cells (Figure 2H; Figures S3A and B), while LDs were rare in this region in cells cultured at glucose replete conditions (data not shown). Interestingly, we found that LDs in proximity to the NVJ displayed prominent areas of tight, evenly spaced contact to the vacuolar membrane. The average distance between vacuoles and the LD surface was 12.4 ± 2.09 nm (mean ± SD) (Figure S3C, left). A prominent feature of these areas were strikingly deformed vacuolar membranes, which appeared sharply bent and curved in a manner allowing for enhanced contact area with multiple adjacent LDs, indicating that the organelles might be physically attached to each other by tether proteins (Figure 2H and Figure S3A, red arrowheads). Furthermore, we frequently observed dense structures at the contact sites that could represent such tethers (Figure 2H, yellow arrowheads). LDs typically form extensive contact sites with the ER membrane (Jacquier et al., 2011), suggesting that accumulation at the NVJ, the interface between the nuclear ER and the vacuole, may reflect simultaneous tethering of LDs to these two partner organelles, and that the LDO proteins might promote formation of LD-vacuole contact sites.

**Figure 2:**
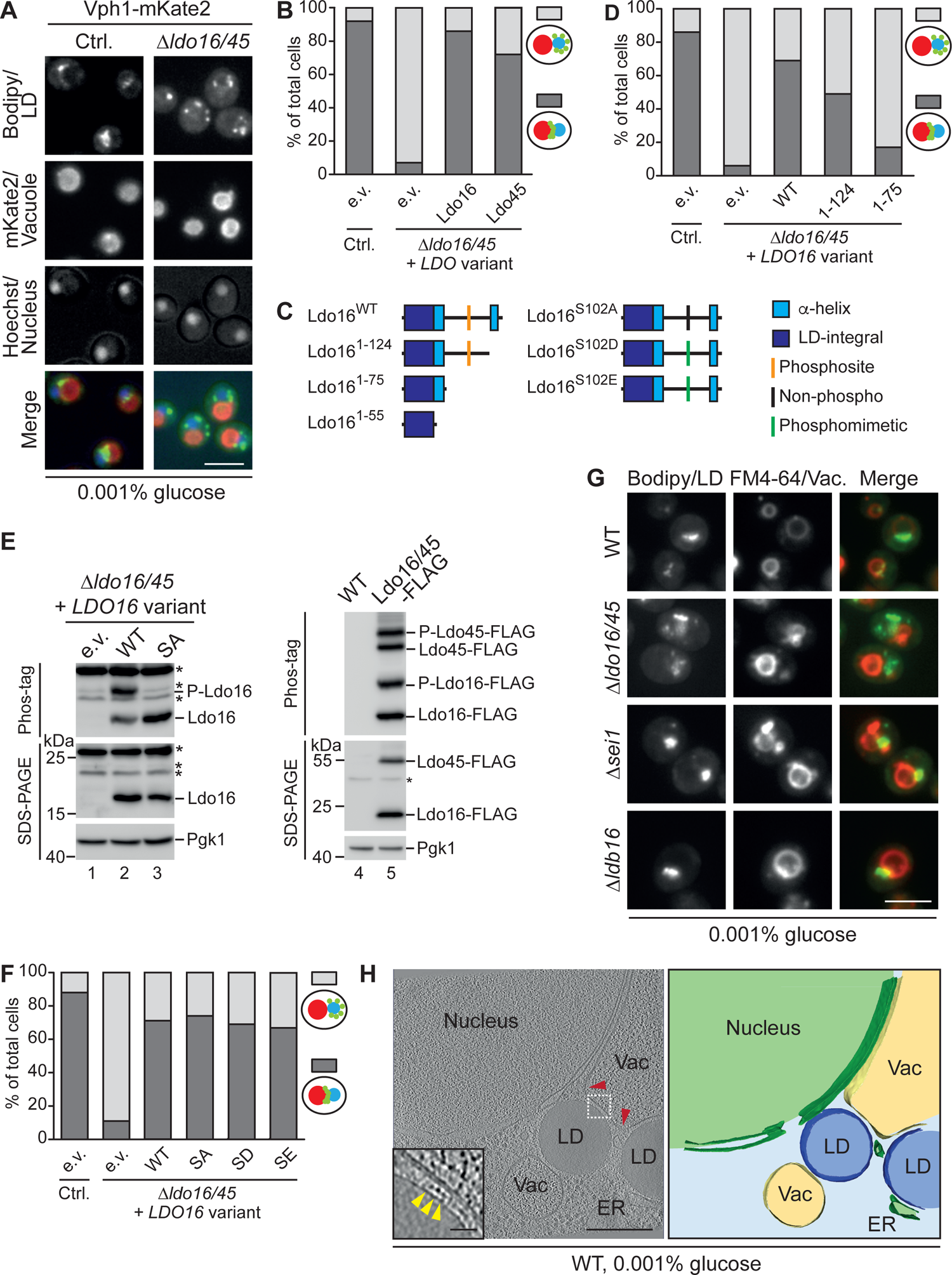
The C-terminal cytosolic domain of Ldo16/45 is required for LD positioning next to the NVJ upon glucose restriction. (A) 1′*ldo16/45* and control cells were cultured overnight in synthetic medium containing 2% glucose, followed by a 4 hour incubation in 0.001% glucose. LDs labeled by the neutral lipid dye BODIPY493/503 (Bodipy) accumulated at the area between the nucleus labeled by Hoechst and the vacuole marked by Vph1-mKate2. Scale bar, 5 µm. (B) Vph1-mKate2 1′*ldo16/45* cells expressing indicated LDO proteins from centromeric plasmids under control of their own promoters were analyzed as described in (A). The number of cells with LDs accumulating at the vacuolar membrane (dark gray) or far from the vacuole (light gray) was quantified. Control (Ctrl.), Vph1-mKate2 cells without *LDO16/45* deletion. n=100 cells. (C) Schematic representation of native Ldo16 (Ldo16^WT^) and of truncated Ldo16 variants. Numbers indicate amino acids present in Ldo16 variants. Secondary structure elements were deleted as indicated. (D) Indicated strains were analyzed as described in (B). n=100 cells. (E) 1′*ldo16/45* cells expressing native Ldo16 (WT) or Ldo16^S102A^ (SA) from centromeric plasmids under control of the *LDO16* promoter (lanes 1-3) or wild-type (WT) cells as well as cells expressing Ldo16/45-FLAG from the genomic LDO locus (lanes 4-5) were cultured to logarithmic growth phase in synthetic medium containing 2% glucose and analyzed by Phos-tag PAGE and SDS-PAGE followed by Western blotting. e.v.: empty vector. Asterisks: non-specific bands. P: phosphorylated forms. (F) 1′*ldo16/45* cells expressing native Ldo16 (WT), non-phospho Ldo16^S102A^ (SA) or phosphomimetic Ldo16^S102D^ (SD) and Ldo16^S102E^ (SE) variants from centromeric plasmids under control of the *LDO16* promoter were analyzed as described in (B). n=100 cells. (G) Wild-type (WT), 1′*ldo16/45,* 1′*sei1,* and 1′*ldb16* cells were cultured overnight in synthetic medium containing 2% glucose, followed by a 4 hour incubation in 0.001% glucose. LDs labeled by the neutral lipid dye BODIPY493/503 (Bodipy) accumulated at the vacuolar membrane labeled by FM4-64 in the absence of the seipin proteins Sei1 and Ldb16. Scale bar, 5 µm. (H) Electron micrograph and tomographic reconstruction of vacuole-LD contact sites in wild-type (WT) cells cultured overnight in synthetic medium containing 2% glucose, followed by a 4 hour incubation in 0.001% glucose. Scale bar, 200 nm. Inset shows electron-dense objects at the LD-vacuole interface that may represent molecular tethers (indicated by yellow arrowheads). Scale bar, 25 nm. 3D segmentation depicted on the right. Vac, vacuole. ER, endoplasmic reticulum.

### The C-terminal cytosolic domain of Ldo16/45 has a role in formation of contact sites between LDs and the vacuole

Our observation of extensive LD-vacuole contacts in glucose restricted wild-type cells suggested that the altered LD distribution in *ldo16/45*1′ cells under these conditions may directly rely on a contact site defect. An LD-vacuole contact site has previously been detected in a systematic study using a bimolecular fluorescence complementation assay, and termed vCLIP for <u>v</u>a<u>c</u>uole <u>lip</u>id droplet contact (Shai et al., 2018), but is unexplored on a molecular and functional level. To test whether Ldo16/45 proteins promote vCLIP formation, we individually overexpressed both proteins in cells cultured at glucose replete conditions, where vCLIPs are rare according to our cryo-electron tomography analyses of wild-type cells. We found that increased LDO levels resulted in an extensive colocalization of LDs and vacuoles, with LDs either forming clusters on vacuolar membranes, or appearing completely enclosed by vacuolar membranes (Figure 3A and Figure S3D). We turned to cryo-electron tomography for high resolution information and found that this colocalization reflected extensive LD-vacuole contact sites (Figure 3B). The distance between LD and vacuole surfaces in these vCLIPs induced by LDO overexpression was 11.7 ± 2.07 nm (mean ± SD), and thus very similar to the dimension of the natural vCLIPs that we had previously observed in wild-type cells during glucose restriction (Figure S3C). Furthermore, we observed vacuolar membrane deformations allowing for enhanced LD contact (Figure 3B; red arrowheads) reminiscent to those observed in natural contact sites. Finally, we detected electron dense putative tethers (Figure 3B; yellow arrowheads) in the synthetically induced vCLIPs similar to those observed in natural vCLIPs.

**Figure 3:**
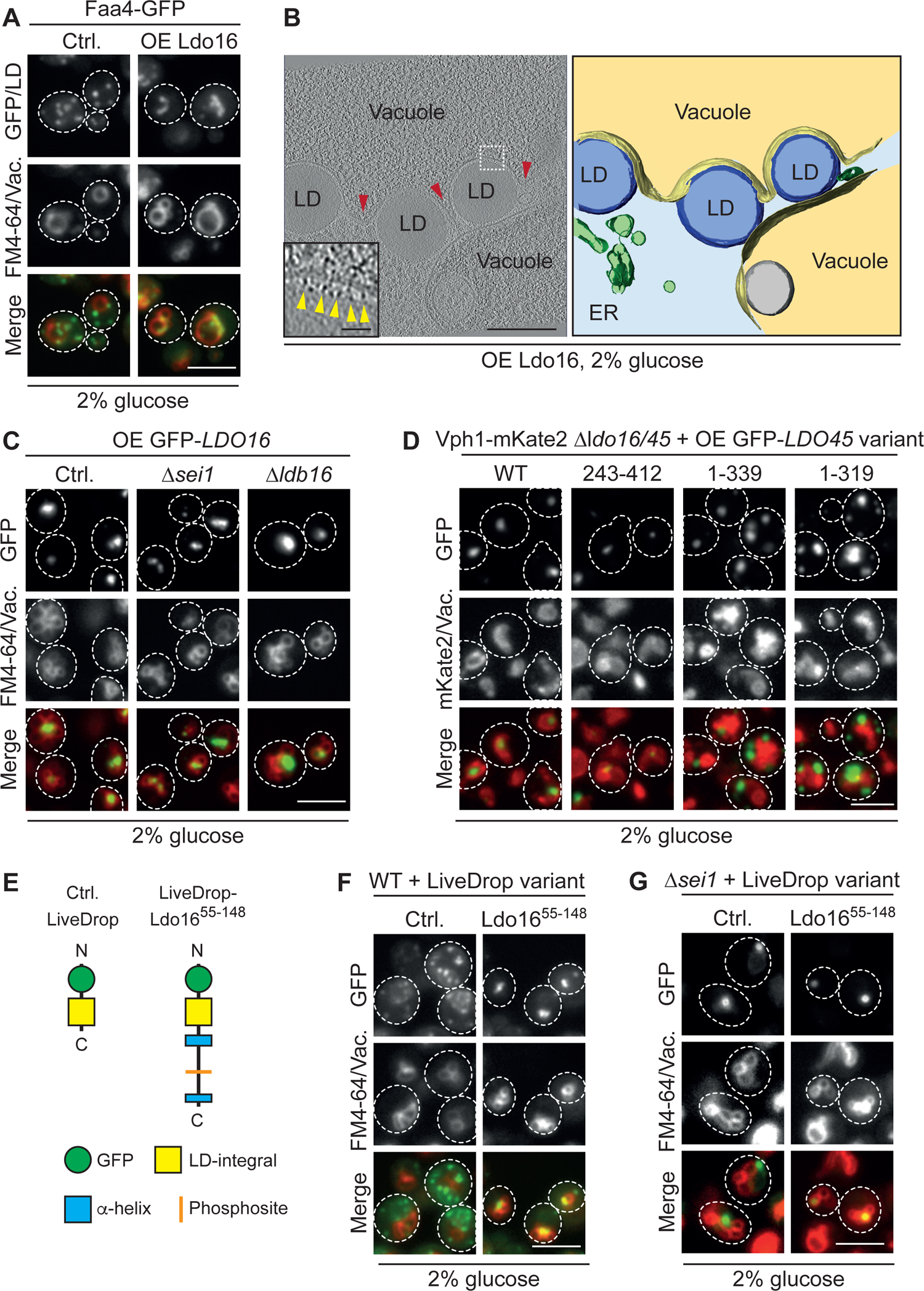
The Ldo16/45 C-terminal soluble domain induces formation of vacuole-LD contact sites (vCLIPs). (A) Cells overexpressing Ldo16 from a *TEF2* promoter were grown to logarithmic phase in synthetic medium containing 2% glucose. LDs were marked by Faa4-GFP, and vacuole membranes were stained with FM4-64. Scale bar, 5 µm. (B) Electron micrograph and tomographic reconstruction from Ldo16 overexpressing strain analyzed in (A). Scale bar, 200 nm. Red arrowheads indicate vacuolar membrane deformations. Inset shows electron-dense objects at the LD-vacuole interface that may represent molecular tethers (indicated by yellow arrowheads). Scale bar, 25 nm. 3D segmentation depicted on the right. ER, endoplasmic reticulum. (C) A centromeric plasmid for overexpression of GFP-Ldo16 from a *TEF2* promoter was transformed into 1′*sei1*, 1′*ldb16*, and control (Ctrl.) cells, which were subsequently grown to logarithmic phase in synthetic medium containing 2% glucose, and stained with FM4-64. Scale bar, 5 µm. (D) Indicated Ldo45 variants were expressed under control of a *TEF2* promoter from centromeric plasmids in cells expressing the vacuole marker Vph1-mKate2. Cells were grown to logarithmic phase in synthetic medium containing 2% glucose, and analyzed by fluorescence microscopy. Scale bar, 5 µm. (E) Schematic representation of standard LiveDrop (Ctrl., control) (left) and LiveDrop-Ldo16^55-148^ (right), a variant carrying the cytosolic C-terminal domain of Ldo16/45. (F) Wild-type (WT) cells expressing LiveDrop variants described in (E) from a centromeric plasmid under control of the *TEF2* promoter were grown to logarithmic growth phase in synthetic medium containing 2% glucose and analyzed by fluorescence microscopy. Scale bar, 5 µm. (G) 1′*sei1* cells were analyzed as described in (F). Scale bar, 5 µm.

Knowing that the colocalization between LDs and vacuolar membranes that we observe by fluorescence microscopy in LDO-overexpressing cells represent extensive vCLIP contact sites (Figures 3A and B; Figure S3D), we used this phenotype as a readout to identify the structural requirements for vCLIP formation. We deleted *SEI1* and *LDB16* in Ldo16-overexpressing cells and found that – similar to natural vCLIPs induced by glucose restriction (Figure 2G) – vCLIPs induced by LDO-overexpression at glucose replete conditions did not require the presence of functional seipin (Figure 3C). Next, we overexpressed truncated variants of Ldo45 in *ldo16/45*1′ cells, to identify the LDO-domains involved in vCLIP formation. We found that deletion of the complete N-terminal cytosolic domain of Ldo45 (variant Ldo45^243-412^) did not affect its ability to induce vCLIPs. In contrast, truncation variants lacking the C-terminal cytosolic domain (Ldo45^1-339^ and Ldo45^1-319^) were unable to induce vCLIPs (Figure 3D). Together with the requirements observed for formation of natural vCLIPs upon glucose restriction (Figure 2D), these findings pointed toward an important role of the C-terminal cytosolic LDO domain in vCLIP formation. We therefore generated a synthetic construct, LiveDrop-Ldo16^55-148^, which consisted of the LD probe LiveDrop fused to the cytosolic LDO-C-terminus (Figure 3E). While overexpression of unmodified LiveDrop had no influence on LD distribution, overexpression of LiveDrop-Ldo16^55-148^ induced LD wrapping in vacuolar membranes (Figure 3F), similar to what we had observed for overexpression of full length LDO proteins (Figure 3A and B). We next expressed LiveDrop and LiveDrop-Ldo16^55-148^ in Δ*sei1* cells. Cells lacking functional seipin had abnormally clustered LDs as expected (Figure 3G). These abnormally shaped LDs were efficiently wrapped by vacuole membranes upon LiveDrop-Ldo16^55-148^ expression (Figure 3G). We conclude that the C-terminus of LDO is both required and sufficient for induction of vCLIP formation, while seipin is not required for this LDO function.

### A microscopy-based screen identifies Vac8 as a player in vCLIP formation

To identify further factors involved in vCLIP formation, we employed an unbiased, systematic approach, where we took advantage of the striking vCLIP phenotype in LDO-overexpressing cells (Figure 3A and B). We used an automated mating approach to introduce an allele for overexpression of GFP-Ldo16 as well as the red vacuole membrane marker Vph1-mCherry into a genome-wide collection of deletion mutants of non-essential genes and hypomorphic alleles of essential genes (Breslow et al., 2008; Giaever et al., 2002) (Figure 4A). This approach resulted in a mutant collection of approximately 6000 strains, which allowed to monitor the effects of loss of function mutations on the formation of Ldo16-dependent vCLIPs. We cultured all mutants at glucose replete conditions, analyzed them by automated microscopy, and searched for mutations that would abolish Ldo16-vacuole colocalization as a measure of vCLIP formation. We identified three classes of hits with an altered relationship between LDs and vacuoles: (1) mutants where the Ldo16-GFP signal was not found in proximity to vacuoles and vCLIPs appeared absent; (2) those where formation of LD clusters typical for Ldo16-overexpressing cells was abolished, while the dispersed Ldo16-GFP signal still colocalized with the vacuolar marker; and (3) a class of ambiguous hits, which displayed diverse mild alterations in the spatial relationship of Ldo16-GFP to the vacuolar marker, or in marker abundance (Figures 4A and B). We re-assessed all hits by staining the vacuole with CMAC and found that in a considerable fraction of the ambiguous class 3 hits, LDs were in fact in proximity to the CMAC positive vacuoles, while the vacuole marker Vph1 appeared mislocalized. This was the case for *atg22* cells and for several ESCRT mutants (*vps4, vps24, vps27, vps2*) (Figure S4A), in which Vph1 localizes to class E compartments (Gerrard et al., 2000; Piper et al., 1995). We reasoned that factors directly involved in the vCLIP tether structure should be found among the class 1 hits of the screen, which comprised only two genes, *VAC8* and *PFA3* (Figure 4B). Vac8 is a multifunctional adaptor protein located on the vacuolar membrane that interacts with multiple partner proteins, including Nvj1 (Pan et al., 2000), Vac17 (Ishikawa et al., 2003; Tang et al., 2003), Atg13 (Scott et al., 2000), and the phosphatidylinositol 3 kinase complex I subunits Vps34 and Vps30 (Lei et al., 2021), together with which Vac8 fulfills a range of biological functions. Interestingly, the second class 1 gene, *PFA3*, encodes a palmitoyltransferase for Vac8 (Hou et al., 2005; Smotrys et al., 2005), and palmitoylation is required for vacuolar localization and full functionality of Vac8 (Subramanian et al., 2006; Wang et al., 1998). As Vac8 mediates formation of NVJ contact sites together with its nuclear ER partner protein Nvj1, and as natural vCLIPs form directly adjacent to NVJs upon glucose starvation, we asked whether the vCLIP defect observed in Δ*vac8* cells was caused by an NVJ defect. To test this, we compared the effect of *VAC8* deletion to that of *NVJ1* deletion, which results in a similar NVJ defect (Pan et al., 2000). However, deletion of *NVJ1* did not alter Ldo16-dependent vCLIPs (Figure 4B), suggesting that Vac8 affects the LD-vacuole contact site independently of NVJ. In a follow-up experiment, we assessed the effects of these mutations on formation of natural vCLIPs upon glucose restriction, and found that it was blocked in Δ*vac8* and Δ*pfa3* cells to an extent comparable to Δ*ldo16* cells, while deletion of *NVJ1* had no effect, again suggesting that Vac8 might have a direct role in vCLIP formation (Figure 4C). To confirm our screening results, we manually constructed a strain overexpressing untagged Ldo16, in which LDs were visualized by expression of the fluorescently labeled LD protein Faa4-GFP, and in which *VAC8* was deleted. While Ldo16-overexpression resulted in colocalization of clustered LDs and vacuolar membranes, absence of Vac8 abolished colocalization but not clustering of LDs (Figure 4D). Vac8 overexpression on the other hand induced vCLIP formation (Figure 4E). We next asked whether Vac8 forms a physical complex with the LDO proteins, as would be expected for components of a tether bridging two organelles. To test this, we performed immunoprecipitation of GFP-Ldo16 and found that Vac8 was indeed specifically co-isolated (Figure 4F). In summary, LDO and Vac8 form a complex and are required for vCLIP formation.

**Figure 4:**
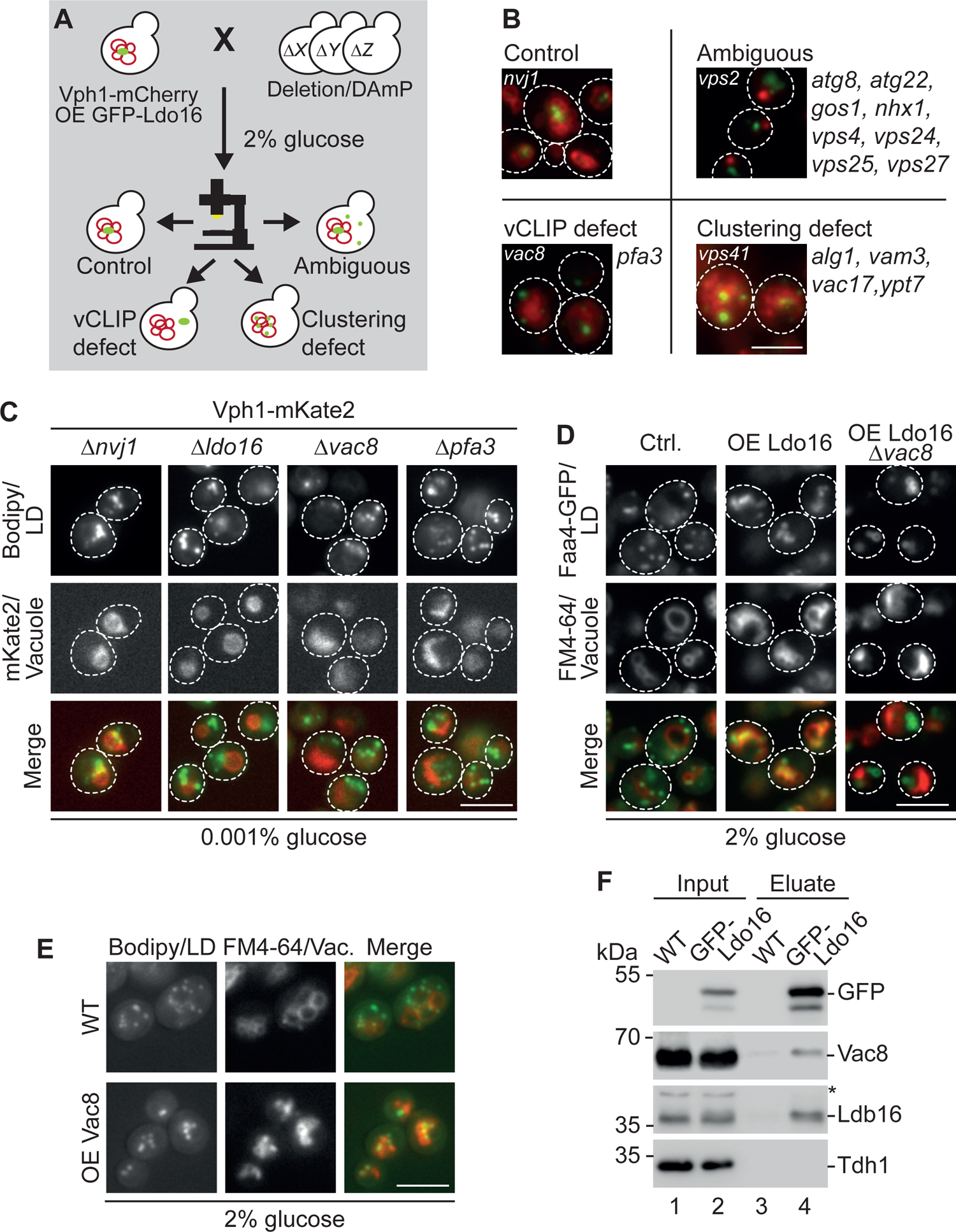
A microscopy-based screen identifies Vac8 as a factor involved in vCLIP formation. (A) Schematic representation of a genome-wide screen searching for genes affecting vCLIP formation upon overexpressing of Ldo16. The vacuolar marker Vph1-mCherry and a p*TEF2-GFP-LDO16* allele were introduced into the genome wide deletion and decreased abundance by mRNA perturbation (DAmP) libraries via automated mating. Mutant strains were grown to logarithmic growth phase in synthetic medium containing 2% glucose, imaged by automated microscopy, and categorized according to indicated traits: Control phenotype with GFP-Ldo16 colocalizing with Vph1-mCherry-labeled vacuole membranes; vCLIP defect, with GFP-Ldo16 and Vph1-mCherry signals not colocalizing; clustering defect, with dispersed GFP signal; and ambiguous, with diverse mild alterations. (B) Representative images of all phenotypic classes described in (A) and lists of all mutations showing the corresponding phenotypes. Scale bar, 5 µm. (C) An allele for expression of Vph1-mKate2 was integrated into indicated deletion strains via automated mating. Before imaging, cells were grown in synthetic medium containing 2% glucose overnight, diluted in medium containing 0.001% glucose, grown for a further 4 hours, and stained with BODIPY493/503 (Bodipy). Strains with vCLIP defects and control cells are shown. Scale bar, 5 µm. (D) An allele for Ldo16 overexpression from a *TEF2* promoter was integrated into cells expressing the LD marker Faa4-GFP and *VAC8* was deleted. Cells were grown to logarithmic phase in synthetic medium containing 2% glucose. Scale bar, 5 µm. (E) Vac8 was overexpressed from a *TEF2* promoter. LDs were stained with BODIPY493/503 (Bodipy) and vacuole membranes with FM4-64. Wild-type (WT) cells were analyzed as negative control. Scale bar, 5 µm. (F) Cells expressing GFP-Ldo16 from a *TEF2* promoter and untagged control cells were subjected to immunoprecipitation using GFP-trap, SDS-PAGE and Western blotting. Input, 5%, Eluate, 100%.

### Ldo16/45 and Vac8 display characteristic features of vCLIP tether proteins

The LDO proteins and Vac8 are required for vCLIP formation upon glucose restriction, and their overexpression results in glucose-independent vCLIP induction. While the LDO proteins are located on the LD surface, and Vac8 at the vacuolar membrane, they are part of the same complex. Based on these collective findings, we hypothesized that an LDO-Vac8 complex could act as a vCLIP tether to physically link the two organelles.

A prominent feature of tether proteins is that they are typically enriched at contact sites. As LDs are too small to reliably resolve LD subregions by conventional fluorescence microscopy, we developed a culture protocol that stimulates vCLIP formation while at the same time promoting LD enlargement. For this, we pre-cultured cells in medium containing 2% glucose, before transferring them into medium lacking glucose, for vCLIP induction, but supplemented with 0.2% oleic acid, which results in enlargement of LDs. Cells expressing Ldo16/45-GFP and the LD marker protein Erg6-mCherry were exposed to this medium for 24 hours, stained with the vacuolar luminal dye CMAC, and analyzed by microscopy. While Erg6-mCherry labeled the complete LD surface in a uniform manner, Ldo16/45-GFP formed bright foci located at the interface between LDs and the vacuole (Figure 5A, left). When we deleted *VAC8*, Ldo16/45-GFP lost its focal localization, and was instead uniformly distributed around the LDs, similar to Erg6-mCherry (Figure 5A, right). We next wanted to analyze Vac8 distribution, and therefore genomically integrated a cassette encoding a fragment for C-terminal GFP-tagging of Vac8 in an Ldo16/45-mCherry background, grew cells in glucose-free, oleic acid-containing medium for 24 hours, and analyzed them by fluorescence microscopy. To our surprise, expression of Vac8-GFP resulted in a complete loss of vCLIPs, with LDs failing to encircle the vacuole and Ldo16/45-mCherry labeling the complete LD surfaces (Figure S5A, left). As a control, we analyzed cells expressing a GFP-tagged variant of the Vac8 partner protein Nvj1, in which Ldo16/45-mCherry formed foci encircling the vacuole as expected (Figure S5A, right). Similarly, LDs accumulated at NVJs labeled by Nvj1-GFP upon glucose starvation, demonstrating efficient vCLIP formation, while this effect was abolished in Vac8-GFP cells (Figure S5B). These findings suggest that similar to *VAC8* deletion, C-terminal tagging of Vac8 blocks vCLIP formation, potentially by abolishing LDO-Vac8 interaction through steric hindrance. This is particularly interesting because *VAC8* deletion mutants have multiple functional defects, which complicates analysis of its specific function in vCLIPs. While vacuoles in Δ*vac8* cells are fragmented (Veit et al., 2001) (Figure 4B-D), vacuoles in Vac8-GFP cells appear morphologically normal (Figure S5A). These cells also efficiently form NVJs (Figure S5A and B), indicating that interaction with the Vac8 partner Nvj1 is unaffected by Vac8 tagging. Expression of Vac8-GFP has also been reported to restore bulk and selective autophagy defects of Δ*vac8* cells (Hollenstein et al., 2019). Collectively, this indicates that Vac8-GFP cells might have a specific vCLIP-defect, while other Vac8 functions remain intact.

**Figure 5:**
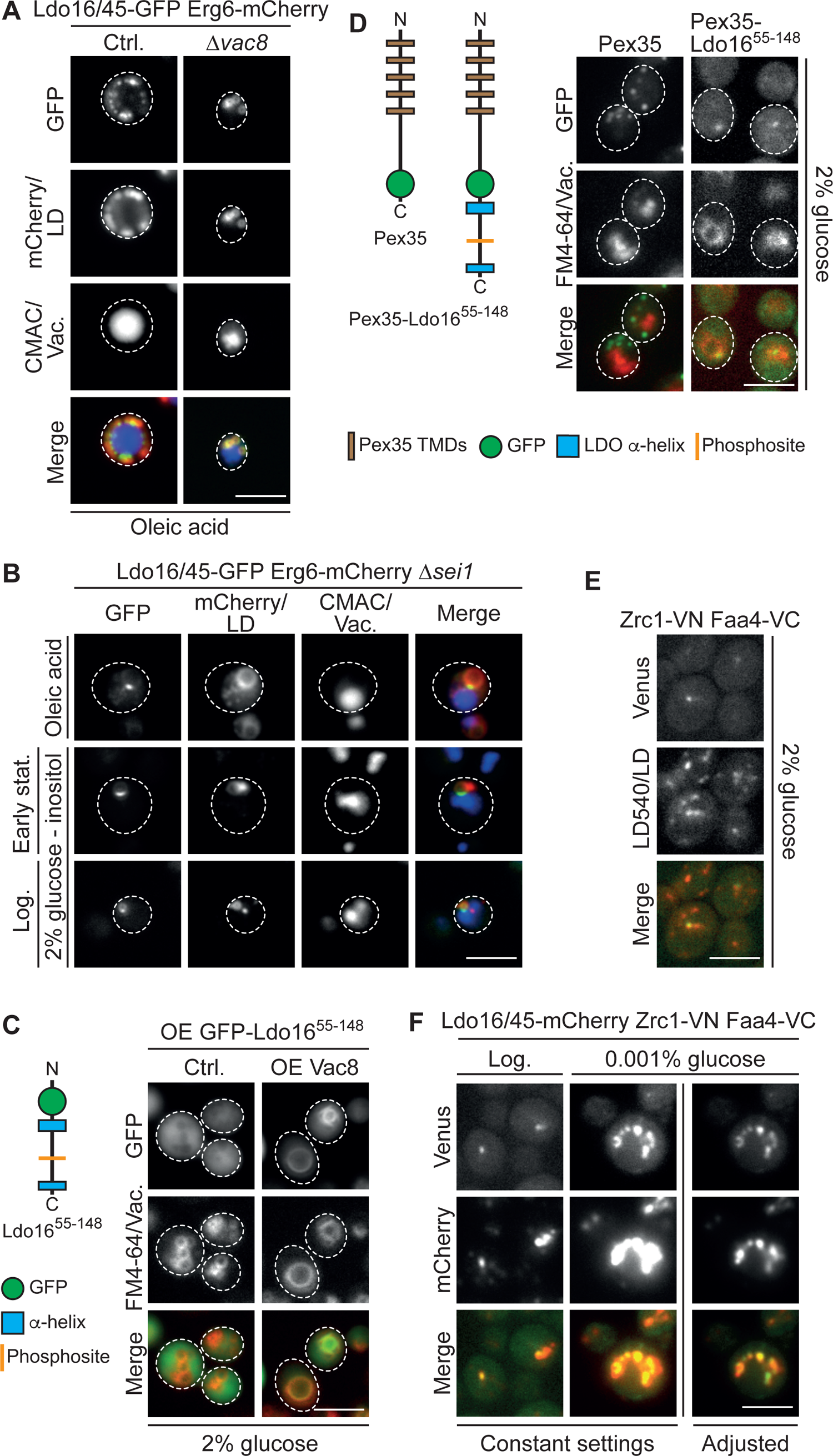
Ldo16/45 and Vac8 are tether proteins at metabolically controlled vCLIPs. (A) Cells expressing Ldo16/45-GFP and Erg6-mCherry with or without deletion of *VAC8* were cultured overnight in synthetic medium containing 2% glucose, and then grown in glucose-free synthetic medium supplemented with 0.2% oleic acid for 24 hours. Cells were stained with the vacuole dye CMAC and analyzed by fluorescence microscopy. Ldo16/45-GFP localized at the interface between LDs and vacuoles in control (Ctrl.) cells, but were uniformly distributed on the LD surface when *VAC8* was deleted. Scale bar, 5 µm. (B) Cells expressing Ldo16/45-GFP and Erg6-mCherry deleted for *SEI1* were grown on inositol-free synthetic medium containing 2% glucose to reach logarithmic phase (bottom) or early stationary phase (middle), or treated with oleic acid as described in (A) (top). Ldo16/45-GFP localized at the interface between LDs and vacuoles (CMAC) in all three conditions. The signal intensity was individually adjusted. Scale bar, 5 µm. (C) Schematic representation of a GFP-tagged Ldo16 variant lacking the LD-anchoring domain (GFP-Ldo16^55-148^) (left). This variant was expressed under control of the *TEF2* promoter from a centromeric plasmid (right). While this Ldo16 variant displayed a cytosolic distribution in control (Ctrl.) cells, overexpression of Vac8 recruited it to the vacuole membrane, stained by FM4-64. Scale bar, 5 µm. (D) Schematic representation of a construct consisting of the peroxisomal membrane protein Pex35, a GFP moiety, and the cytosolic domain of Ldo16, as well as a Pex35-GFP control construct (left). These proteins were expressed under a *TEF2* promoter from a centromeric plasmid. Cells were grown to logarithmic growth phase in synthetic medium containing 2% glucose, stained with the vacuolar membrane dye FM4-64, and analyzed by fluorescence microscopy. Scale bar, 5 µm. (E) Non-fluorescent halves of Venus were fused to the vacuolar membrane protein Zrc1 (Zrc1-VN) and the LD protein Faa4 (Faa4-VC). Cells were grown to logarithmic growth phase in synthetic medium containing 2% glucose, and LDs were stained by LD540. A subpopulation of LDs close to the vacuole was labeled by bimolecular fluorescence complementation, reporting on a vCLIP contact site. Scale bar, 5 µm. (F) Vacuole-associated LDs were detected by bimolecular fluorescence complementation as described in (E) in cells expressing Ldo16/45-mCherry. Cells were grown to logarithmic growth phase in synthetic medium containing 2% glucose (left) or transferred for 4 hours to medium containing 0.001% glucose (middle and right). Right column shows the same field of view as middle column, with adjusted signal intensity. Scale bar, 5 µm.

We next deleted *SEI1* to test whether seipin is required for positioning of Ldo16/45 at vCLIPs. We found that exposure to oleic acid resulted in formation of enlarged LDs in the Δ*sei1* mutant that were consistently found adjacent to vacuoles, and Ldo16/45-GFP formed bright foci at the organelle interfaces. Δ*sei1* cells also form supersized LDs in response to inositol depletion (Wang et al., 2014b), and LDO proteins accumulated efficiently at vCLIPs independently of seipin also under this condition (Figure 5B).

We next overexpressed a fluorescently tagged Ldo16 variant lacking the hydrophobic, LD-integral part, GFP-Ldo16^55-148^ (Figure 5C). We reasoned that if LDO proteins and Vac8 form a vCLIP tether complex, then this Ldo16 variant should localize to the vacuole in a manner dependent on Vac8. When expressed in control cells, GFP-Ldo16^55-148^ showed a cytosolic pattern (Figure 5C), confirming that the hydrophobic N-terminal part of the protein (residues 1-55) is important for LD binding. In contrast, when we overexpressed Vac8, GFP-Ldo16^55-148^ uniformly accumulated on the vacuolar surface (Figure 5C), distinct from the LD clustering patterns observed previously (Fig. 4E). This shows that the C-terminal cytosolic domain of LDO proteins has the structural capacity to bind to the vacuolar membrane, in a manner dependent on Vac8.

To ultimately clarify whether the LDO proteins act as molecular tether proteins, we designed an experiment to test whether they can synthetically recruit other cellular structures to the vacuolar membrane. For this, we generated a construct Pex35-GFP-Ldo16^55-148^ consisting of the peroxisomal membrane protein Pex35 (Yofe et al., 2017), followed by a GFP tag for visualization, and finally the cytosolic domain of Ldo16 at the very C-terminus (Figure 5D). As a control, we used a similar Pex35-GFP construct lacking the Ldo16 domain. Both Pex35-GFP-Ldo16^55-148^ and Pex35-GFP colocalized with an RFP-PTS1 marker, indicating that they were efficiently targeted to peroxisomes (Figure S5C). When we labeled vacuolar membranes using FM4-64, we found that overexpression of Pex35-GFP-Ldo16^55-148^ resulted in colocalization of peroxisomes with vacuolar membranes, while vacuoles were far away from peroxisomes in the control cells (Figure 5D). This demonstrates that the LDO C-terminus, when attached to the surface of peroxisome, is sufficient to induce synthetic peroxisome-vacuole contact sites.

Collectively, these results show that the LDO proteins act as contact site tether proteins at the vCLIP, where they attach LDs to vacuoles in a manner dependent on the vacuolar surface protein Vac8.

### vCLIP abundance is regulated by the metabolic state of the cell

We have reported in the past that the LDO proteins are strongly enriched on a subpopulation of LDs in cells during logarithmic growth phase at glucose replete conditions (Eisenberg-Bord et al., 2018). In light of our finding that LDO is involved in vCLIP formation upon glucose restriction, we hypothesized that the LDO-enriched LDs observed in logarithmic phase could represent a specialized LD subpopulation engaged in a vCLIP even at glucose replete conditions. During logarithmic phase, multiple small LDs are typically present in each cell, and reliably monitoring the exact spatial relationship of an LD subpopulation to the vacuole by conventional fluorescence microscopy is challenging. We therefore turned to a contact site reporter technique based on the split Venus system (Eisenberg-Bord et al., 2016). We fused one half of split Venus to the LD surface protein Faa4, and the other half to the vacuolar membrane protein Zrc1. This pair of split Venus fusion constructs acts as a vCLIP contact site reporter that allows to detect close proximity between the organelle surfaces by spatially restricted fluorescence complementation. We stained LDs in logarithmic phase cells expressing the vCLIP reporter with the neutral lipid dye LD540 and found that indeed, just one or two LDs per cell were vCLIP reporter positive (Figure 5E). We then introduced the vCLIP reporter into a strain expressing Ldo16/45-mCherry. The contact site reporter co-localized with Ldo16/45-mCherry (Figure 5F), demonstrating the presence of an LD subpopulation engaged in Ldo16/45-mediated vCLIPs during logarithmic growth phase. We then analyzed the same strain at glucose restriction. At this condition, both the vCLIP reporter and Ldo16/45-mCherry labeled numerous LDs, showing that the LD subpopulation engaged in vCLIPs expands strobgly in response to glucose restriction (Figure 5F).

### Ldo45 combines LD-vacuole tethering- and protein-recruitment activities, and determines localization of Pdr16 at vCLIP

The notion that the LDO proteins and Vac8 are responsible for formation of vCLIPs raised the question how Pdr16, which requires Ldo45 for targeting to LDs (Figure 1) (Eisenberg-Bord et al., 2018; Teixeira et al., 2018), is related to this contact site. We introduced the split Venus-based vCLIP reporter into a Pdr16-mCherry strain and grew the cells at glucose-replete conditions to logarithmic phase. Similar to what we had previously seen for Ldo16/45 (Figure 5F), Pdr16-mCherry localized specifically to the vCLIP-positive LD subpopulation (Figure 6A). To test whether Pdr16 localizes to the complete surface of these LDs, or to the vacuole-LD interface itself, we grew Pdr16-GFP Erg6-mCherry cells on 0.2% oleic acid, and stained the vacuoles with CMAC. We found that similar to the LDO proteins (Figure 5A), Pdr16 formed foci at the sites where LDs were apposed to the vacuolar surface (Figure 6B), suggesting that Ldo45 recruits Pdr16 to vCLIPs.

**Figure 6:**
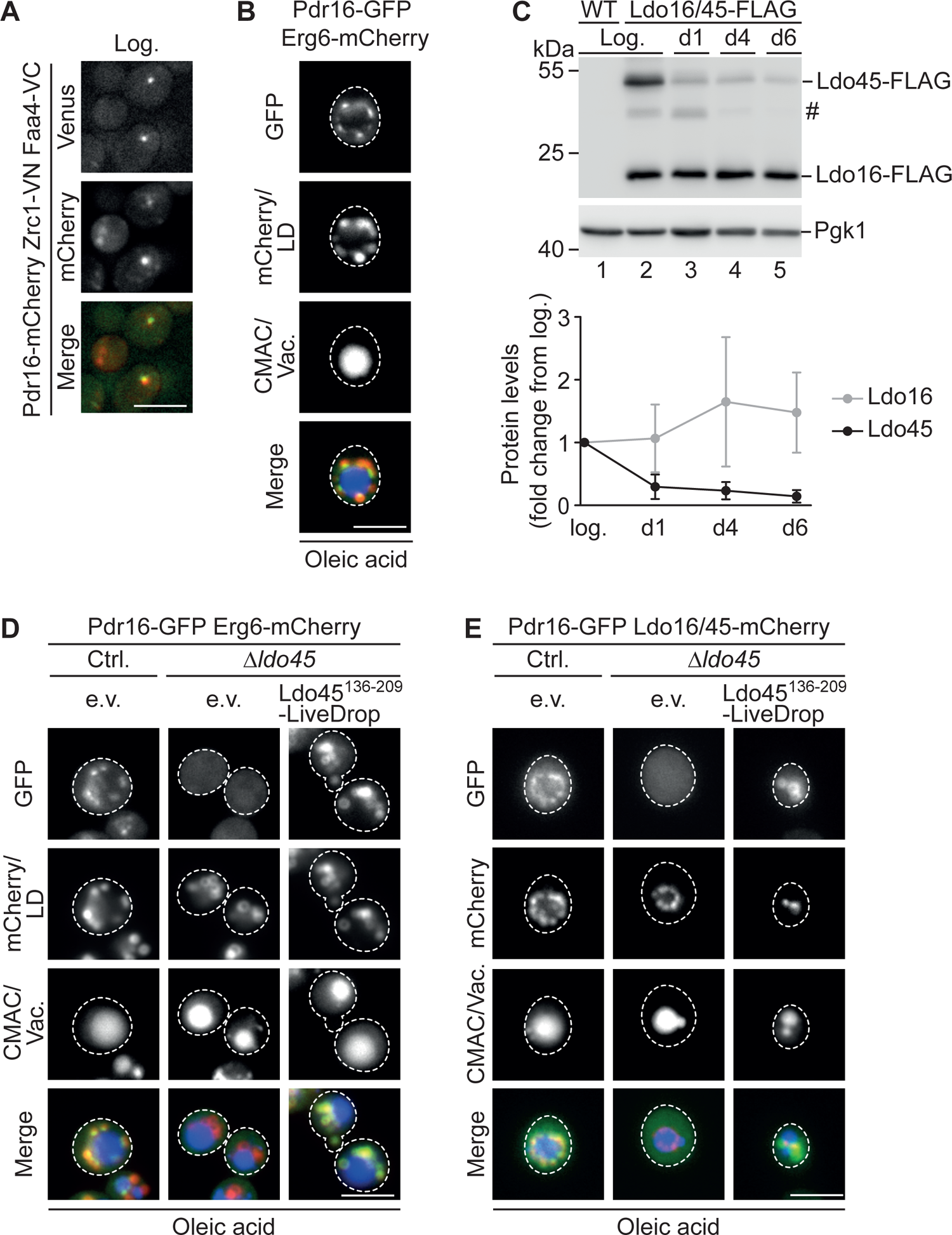
Structural coupling of LD-tethering and Pdr16-recruitment activities of Ldo45 mediates Pdr16 accumulation at vCLIP. (A) Non-fluorescent halves of Venus were fused to the vacuolar membrane protein Zrc1 (Zrc1-VN) and the LD protein Faa4 (Faa4-VC). Pdr16-mCherry colocalized with the vacuole-associated LD subpopulation visualized by bimolecular fluorescence complementation in cells grown to logarithmic growth phase in synthetic medium containing 2% glucose. Scale bar, 5 µm. (B) Pdr16-GFP Erg6-mCherry cells were grown in synthetic medium supplemented with 0.2% oleic acid for 24 hours. Pdr16-GFP formed foci at the interface of LDs (Erg6-mCherry) and vacuoles (CMAC). Scale bar, 5 µm. (C) Top: Ldo16/45-FLAG and wild-type (WT) cells were grown in in synthetic medium containing 2% glucose and lysates were extracted at indicated states: logarithmic growth phase (log.), and 1 day (d1), 4 days (d4), and 6 days (d6) growth into stationary growth phase. Samples were analyzed by SDS-PAGE and Western blotting. #, band of unclear identity. Bottom: Protein levels of Ldo16-FLAG and Ldo45-FLAG normalized to the Pgk1 loading control are shown as fold change from logarithmic growth phase. n=3. Error bars indicate SD values. (D) Pdr16-GFP Erg6-mCherry cells with or without *LDO45* deletion were grown in synthetic medium supplemented with 0.2% oleic acid for 24 hours and stained with the vacuole dye CMAC. While Pdr16-GFP accumulated at interfaces between LDs (Erg6-mCherry) and vacuoles (CMAC) in control (Ctrl.) cells, deletion of *LDO45* resulted in a cytosolic Pdr16-GFP distribution. Expression of Ldo45^136-209^-LiveDrop from a centromeric plasmid under control of a *TEF2* promoter resulted in recruitment of Pdr16-GFP to LDs, but failed to focus Pdr16-GFP at vCLIPs. e.v., empty vector. Scale bar, 5 µm. (E) The experiment was performed as described in (D), with the difference that Ldo16/45-mCherry for vCLIP labeling was expressed instead of the LD surface marker Erg6-mCherry. Scale bar, 5 µm.

Ldo45 levels have been reported to be metabolically controlled, and to drop when cells enter stationary growth phase (Teixeira et al., 2018). We analyzed Ldo16/45-FLAG cells in different growth phases by Western blotting. Consistent with the findings by Teixeira et al. (2018), we detected a drop in Ldo45 levels in stationary phase, while Ldo16 levels were stable (Figure 6C). Nonetheless, we detected some Ldo45 even in late stationary phase (Figure 6C). As the *LDO16* and *LDO45* genes overlap with each other (Figure S1A), introduction of a tagging cassette downstream to the locus results in simultaneous tagging of Ldo16 and Ldo45, so that it was unclear whether our Ldo16/45-GFP strain, which labels the LD-vacuole interface (Figure 5A), reports on localization of both or just one LDO variant at the vCLIP. We therefore used a strain expressing from its genomic locus N-terminally tagged GFP-Ldo45 under control of its endogenous promoter (Yofe et al., 2016), and introduced a plasmid for expression of Ldo16-mCherry from its endogenous promoter. We supplemented the strain with oleic acid to induce LD enlargement, and analyzed the cells by CMAC labeling and fluorescence microscopy. We found that both GFP-Ldo45 and Ldo16-mCherry co-localized at LD-vacuole interfaces (Figure S6), suggesting that two alternative vCLIP tethers, Ldo16-Vac8 and Ldo45-Vac8, exist.

Our structure function analysis has shown that the C-terminal cytosolic domain present in both Ldo16 and Ldo45 mediates Vac8-dependent tethering (Figure 2 and 3), while Pdr16 targeting to LDs depends on an amphipathic helix in the N-terminal cytosolic domain that is only present in Ldo45 (Figure 1). We asked whether direct coupling of the C-terminal tether-domain and the N-terminal PITP targeting-domain in one protein, Ldo45, is a prerequisite for Pdr16 localization to the organelle interface. We co-expressed Pdr16-GFP either with Erg6-mCherry (Figure 6D) or with Ldo16/45-mCherry (Figure 6E), and analyzed cells by fluorescence microscopy upon supplementation with oleic acid, enabling visualization of Pdr16 foci together with Ldo16/45 at LD-vacuole interfaces. We then deleted *LDO45*, which results in loss of Ldo45, while Ldo16 expression is maintained (Figure 1A and Figure S1A). In these cells, Ldo16-mCherry was still enriched at the vCLIP, while Pdr16 was dispersed in the cytosol as expected (Figure 6D and E). We then introduced a plasmid Ldo45^136-209^-LiveDrop, comprising the Ldo45 amphipathic that we found to mediate Pdr16 targeting to LDs (Figure 1). In these cells, Pdr16 was efficiently targeted to LDs, but failed to concentrate at the LD-vacuole interface labeled by Ldo16-mCherry, and showed instead a uniform distribution across the LD surface (Figure 6D and E).

We conclude that vCLIP depends on two structurally distinct tethers: One consists of Ldo16 and Vac8 and mediates organelle tethering, but has no relationship to Pdr16. The other tether consists of Vac8 and Ldo45, which, in addition to its tethering function, recruits the PITP Pdr16 to the contact site.

### The tethering function of vCLIP has a role in LD breakdown by lipophagy

Cells lacking the LDO proteins have previously been reported to display a lipophagy defect (Teixeira et al., 2018). To monitor this process, we grew cells expressing the vacuolar membrane marker Vph1-mKate2 to late stationary growth phase and stained LDs with Bodipy, allowing for visualization of LDs in the cytosol, as well as those internalized into the vacuole by lipophagy. We found that absence of LDO proteins resulted in a lipophagy block, as reported (Figure 7A and B). We re-expressed either Ldo16 or Ldo45 individually from plasmids from their own promoters, and found that either LDO variant efficiently supported lipophagy (Figure 7A and B). Of note, plasmid-born Ldo45 failed to rescue lipophagy in an *ldo* deletion mutant in a previous study (Teixeira et al., 2018), a discrepancy that may be caused by differences in Ldo45 expression levels. To understand the role of LDO proteins in lipophagy, we next tested whether the LDO partner proteins Pdr16 or Vac8 were required for this process. Lipophagy occurred normally in Δ*pdr16* cells (Figure 7C and Figure S7A). This is consistent with the observation that Ldo16, which cannot recruit Pdr16, supports lipophagy in the absence of Ldo45. Δ*vac8* mutants had a block in lipophagy similar to 1′*ldo16/45* mutants (Figure S7A). A lipophagy defect in the absence of *VAC8* has also been reported in a previous study (van Zutphen et al., 2014). Due to the multiple roles of Vac8, Δ*vac8* cells have a range of defects, including an abnormal vacuolar morphology. To test whether the lipophagy defect of Δ*vac8* cells was directly linked to vCLIP, we assessed cells expressing the Vac8 variant Vac8-GFP, which fails to form vCLIPs similar to Δ*vac8* cells, but has vacuoles with a normal morphology (Figure S5A). Similar to Δ*vac8* cells, Vac8-GFP cells displayed a lipophagy defect (Figure 7C and Figure S7A). Together, these findings suggest that vCLIP-mediated LD-vacuole tethering is important for the process of lipophagy. We next analyzed the ability of Ldo16 truncation variants to support lipophagy, and found that their ability to support LD accumulation on the vacuolar surface (Figure 2D) and the ability to support lipophagy correlated, with Ldo16^1-124^ showing a partial lipophagy defect, while Ldo16^1-75^ was virtually unable to mediate LD internalization into the vacuole (Figure 7D and Figure S7B). In addition, we also assessed lipophagy in cells expressing the phosphomimetic variants Ldo16^S102D^ and Ldo16^S102E^, as well as the non-phosphorylatable mutant Ldo16^S102A^. While the S102A variant supported lipophagy similar to wild-type Ldo16, the phosphomimetic variants surprisingly both had a strong lipophagy defect (Figure 7E and Figure S7C). We tested protein levels of all Ldo16 phosphosite variants in late stationary phase, during lipophagy, and found that the proteins were stable (Figure S7D). We therefore monitored phosphorylation of native Ldo16 by Phos-tag SDS PAGE throughout the protocol for stationary phase-induced lipophagy. We found that Ldo16 was phosphorylated in cells in logarithmic growth phase, and became gradually dephosphorylated when cells progressed toward late stationary phase, when lipophagy occurs (Figure 7F). This suggests that the ability of Ldo16 to efficiently support lipophagy requires dephosphorylation at position serine 102.

**Figure 7:**
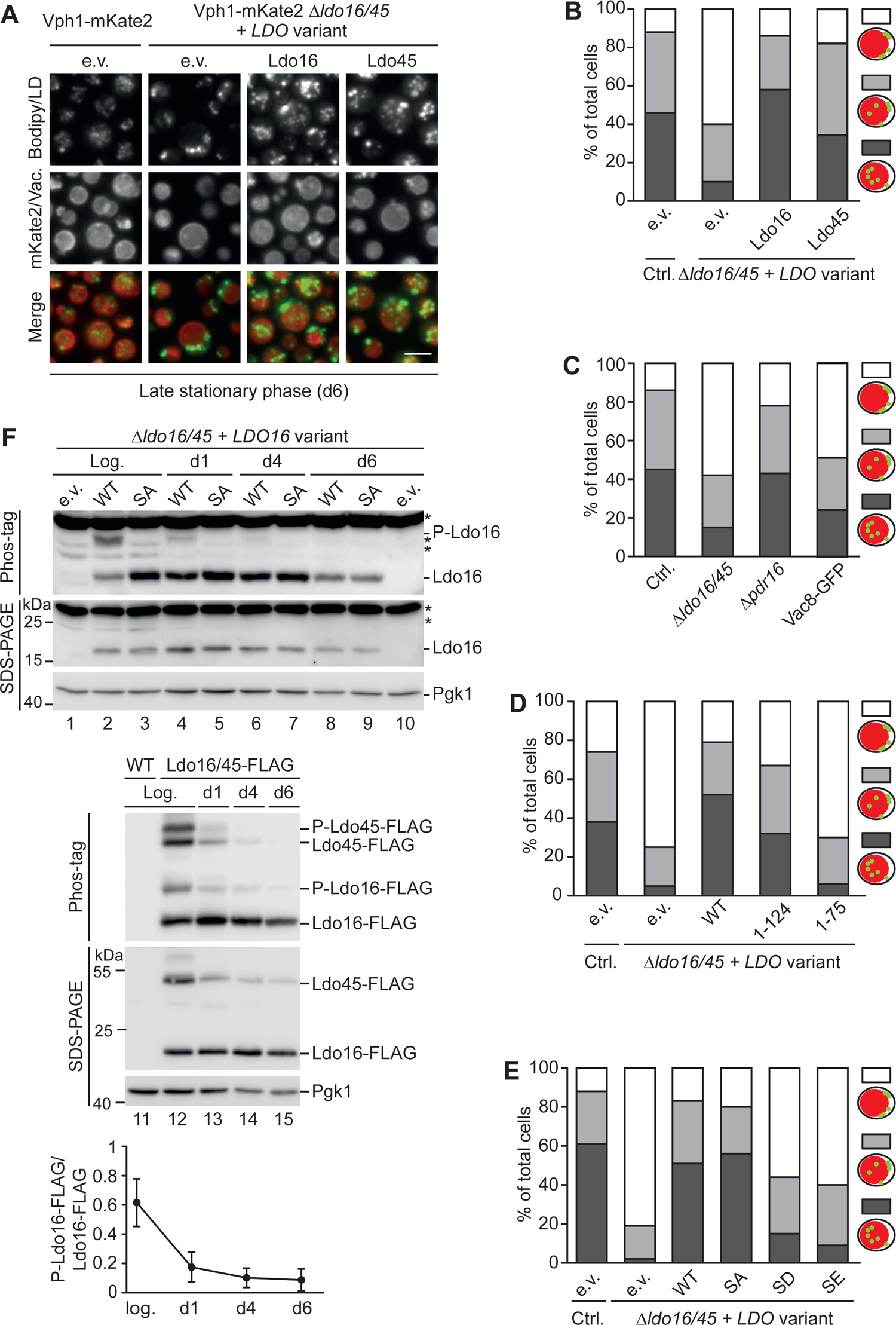
Ldo16/45 phosphorylation state affects internalization of vacuole-tethered LDs by lipophagy. (A) Indicated strains were cultured overnight in synthetic medium supplemented with 2% glucose, diluted in the same medium, and grown for 6 days (d6) into late stationary phase to induce lipophagy. Bodipy-stained LDs were internalized into the vacuole labeled by Vph1-mKate2 by lipophagy, which was abolished in the absence of Ldo16/45. Centromeric plasmids individually encoding Ldo16 or Ldo45 under the control of their own promoters rescued this lipophagy defect. e.v., empty vector. Scale bar, 5 µm. (B) Cells in (A) were classified into three groups according to the number of LDs located inside the vacuolar lumen: no LDs inside the vacuole (white), 1-3 LDs inside the vacuole (light gray), >3 LDs inside the vacuole (dark gray). n=50 cells. (C) Control (Ctrl.), 1′*ldo16/45*, 1′*pdr16*, and Vac8-GFP cells were analyzed as described in (A) and results were quantified as described in (B). n=100 cells. (D-E) 1′*ldo16/45* cells expressing indicated Ldo16 variants from centromeric plasmids under control of the *LDO16* promoter or empty vector (e.v.) controls were analyzed as described in (A) and results were quantified as described in (B). n=100 cells. (F) Top, lanes 1-10: 1′*ldo16/45* cells expressing native Ldo16 (WT) or Ldo16^S102A^ (SA) from centromeric plasmids under control of the *LDO16* promoter were cultured in synthetic medium containing 2% glucose and harvested at different time points: logarithmic growth phase (log.), or day 1 (d1), day 4 (d4), day 6 (d6) of growth into stationary phase. Proteins were extracted and analyzed by Phos-tag PAGE or standard SDS-PAGE and Western blotting. Middle, lane 11-15: Cells expressing Ldo16/45-FLAG from the genomic *LDO* locus and wild-type (WT) controls were analyzed in the same way. P marks phosphorylated LDO species. Bottom: The ratio between phosphorylated and dephosphorylated Ldo16-FLAG species was quantified. n=3. Error bars indicate SD values.

In summary, our results suggest an involvement of vCLIP in lipophagy. LD-vacuole tethering, but not Pdr16 recruitment, are required for LD internalization into the vacuole, a process that might be regulated by the LDO phosphorylation status.

## DISCUSSION

We have identified a critical protein machinery of the vCLIP, a poorly characterized contact site between LDs and the vacuole. vCLIPs are formed by at least two types of structurally closely related molecular tethers, which consist of the armadillo-repeat protein Vac8 on the vacuolar side in complex with either Ldo16 or Ldo45 on the LD side. Ldo45, but not Ldo16, is additionally linked to the PITP Pdr16, and responsible for recruitment of this protein to vCLIP.

Vac8 is a multifunctional protein that has roles in vacuole inheritance (Pan and Goldfarb, 1998; Wang et al., 1996, 1998), in vacuolar fusion (Pan and Goldfarb, 1998; Veit et al., 2001), in bulk-(Gatica et al., 2021; Hollenstein et al., 2019) as well as different forms of selective autophagy (Boutouja et al., 2019; Kissová et al., 2007; Oku et al., 2006; van Zutphen et al., 2014), in the CVT pathway (Wang et al., 1996, 1998), and in formation of the NVJ contact site (Elbaz-Alon et al., 2015; Murley et al., 2015; Pan et al., 2000). The LDO proteins on the other hand have previously been identified by us and others as partner proteins of the LD biogenesis machinery seipin, which acts at LD-ER contact sites (Eisenberg-Bord et al., 2018; Teixeira et al., 2018). We find that LDO-dependent vCLIP formation occurs efficiently in the absence of functional seipin, indicating that LDOs fulfill a dual role, on one hand at LD-ER interfaces, together with seipin, and on the other hand at LD-vacuole interfaces, together with Vac8 and Pdr16. Multifunctional proteins with distinct interaction partners like Vac8 and Ldo16/45 are perfectly suited for coordinating cellular responses to metabolic cues, by acting as molecular switches that control different biological processes in an orchestrated manner. Consistently, the abundance of vCLIPs responds strongly to the metabolic state of the cell. While only a small subpopulation of LDs, typically 1-2 per cell, are tethered to the vacuole in logarithmic growth phase, the vCLIP-positive LD pool expands drastically once cells are exposed to nutrient stress, such as glucose restriction, or growth on oleate. In addition to the regulation of vCLIP abundance, also vCLIP composition might adapt to metabolic cues. In agreement with observations from Teixeira et al., (2018), we observe that while Ldo16 levels are stable across different metabolic states, Ldo45 levels are regulated, and decrease during progression into stationary growth phase, when cells run out of glucose. Ldo45 in turn is required for recruitment of Pdr16 to vCLIPs, indicating that Pdr16 abundance at the contact site might be tightly temporally controlled, and that the vCLIP proteome and thus likely also function might be altered when cells progress through stationary phase. Pdr16 is part of a family of Sec14-like PITPs (van den Hazel et al., 1999; Li et al., 2000). Sec14-family proteins comprise hydrophobic cavities that can accommodate single lipid molecules (Sha et al., 1998). In vitro, Pdr16 shows phosphatidylinositol transfer activity (Li et al., 2000). Sec14-family proteins typically bind a second type of lipid besides phosphatidylinositol, which in the case of Pdr16 has been suggested to be a sterol (Tripathi et al., 2019), potentially lanosterol (Šťastný et al., 2022). Besides structural tether proteins, contact sites typically comprise functional proteins specialized for material transport between the tethered membranes, and it is conceivable that Pdr16 could mediate transfer of phosphatidylinositol and sterols between LDs and the vacuole. LDs store large amounts of sterols in their central core compartment, in the form of sterol esters. Furthermore, in stationary growth phase, when vCLIPs form extensively, vacuolar membranes segregate into liquid disordered and sterol-rich, liquid ordered domains (Toulmay and Prinz, 2013), suggesting that sterol exchange could indeed be a vCLIP function. However, the role of Sec14-family members as lipid transfer proteins in vivo is a matter of debate. Instead, these proteins have been reported to fulfill regulatory roles in phosphoinositide signaling, by stimulating specific lipid metabolism enzymes (Schaaf et al., 2008).

Pdr16 has been shown to counteract lipolytic LD breakdown during meiosis in a manner dependent on phosphatidylinositol 4-phosphate (Ren et al., 2014). In yeast, meiosis takes place when diploid cells are subjected to severe nutrient deprivation. This indicates that Pdr16 might limit lipolytic LD breakdown when lipids become more precious due to restrictions in nutrient availability, a function that may also apply for vegetative haploid cells exposed to glucose restriction. Consistently, Bsc2, a protein that we previously identified as a further resident of the LDO- and Pdr16-enriched LD subpopulation of cells in logarithmic growth phase (Eisenberg-Bord et al., 2018), was recently shown to counteract triacylglycerol lipolysis (Speer et al., 2023), suggesting that the LD subpopulation engaged in vCLIP during logarithmic phase might be protected from lipolysis. Similar to what we observe for metabolic regulation of vCLIP abundance, the pool of Bsc2-positive LDs expands when cells run out of glucose (Speer et al., 2023). Collectively, this suggests that LDs engaged in vCLIPs may be preserved for longer-term lipid storage particularly during nutrient deprivation.

Prolonged nutrient restriction ultimately leads to LD degradation by lipophagy, allowing for lipid mobilization and utilization as carbon source (Seo et al., 2017; Tsuji et al., 2017; Vevea et al., 2015; Wang et al., 2014a; van Zutphen et al., 2014). We find that vCLIP-mediated LD-vacuole tethering is required for this process in late stationary phase. Lipophagy is largely blocked in cells lacking the vCLIP tether proteins Ldo16/45 and Vac8, and also hampered in mutants expressing a GFP-tagged Vac8 variant, in which vCLIPs are defective, while other Vac8 functions are maintained. Pdr16 on the other hand is not required for lipophagy. The C-terminal tether domain of Ldo16 and presumably also that of Ldo45 is phosphorylated (on serine 102 of Ldo16) in logarithmic phase, but dephosphorylated in stationary phase. Serine 102 is a proline-directed site and thus likely phosphorylated by a member of the CMGC kinase family, but the specific kinase and phosphatase responsible for controlling the Ldo16 phosphorylation state are currently unknown. While a non-phosphorylatable Ldo16 variant efficiently supports lipophagy in late stationary phase, phosphomimetic Ldo16 mutants have a lipophagy defect. This suggests that information on the metabolic state of the cell may be integrated through LDO (de-)phosphorylation, and determine the fate of vacuole-tethered LDs upon activation of the lipophagic machinery.

LDAF1/promethin proteins are metazoan proteins with similarity to LDOs. These proteins also form a complex with seipin (Castro et al., 2019; Chartschenko et al., 2021; Chung et al., 2019), and their loss affects the process of LD biogenesis (Chung et al., 2019). It remains to be determined to which extent the multifunctional character that we observe for the yeast LDO proteins is conserved to metazoans, and whether these proteins have a role in lipid droplet turnover and/or contact sites.

More broadly, we have described the molecular machinery underlying formation of a metabolically controlled organelle contact site, vCLIP, which has a central role in controlling LD turnover. As the vCLIP tether proteins Ldo16 and Ldo45 are partner proteins of the LD biogenesis machinery seipin, these findings imply a direct molecular link between the processes of LD formation and consumption, and a central role of the LDO proteins in coordinating lipid homeostasis.

## ACKNOWLEDGEMENTS

This work was supported by the Gerty Cori Programme, Medical Faculty, University of Münster (to M.B.), and by the Deutsche Forschungsgemeinschaft (DFG, German Research Foundation), SFB1190 P21 (to M.B.) and P22 (to R.F.-B). J.C. and R.F.-B. acknowledge funding from Germany’s Excellence Strategy (EXC 2067/1-390729940). Cryo-ET instrumentation at the University of Göttingen was jointly funded by the DFG Major Research Instrumentation program (448415290) and the Ministry of Science and Culture of the State of Lower Saxony. O.S. is supported by the Austrian Science Fund (FWF) [P36187-B]. D.T.V.D. and M.H. are members of CiM-IMPRS, the joint graduate school of the Cells-in-Motion Interfaculty Centre, University of Münster, Germany and the International Max Planck Research School – Molecular Biomedicine, Münster, Germany. We thank Christoph Thiele for the LD540 neutral lipid dye; Maya Schuldiner, Roland Wedlich-Söldner, Naama Barkai, Tomer Ravid and Jeffrey Gerst for plasmids; Christian Ungermann for the Vac8 antibody; Sanjana Rao for embarking on the Phos-tag adventure; Tat Cheng for electron microscopy support; and Roland Wedlich-Söldner, Christian Schuberth, Maya Schuldiner, Einat Zalckvar, Michal Eisenberg-Bord, and all members of the Bohnert lab for discussion.

## AUTHOR CONTRIBUTIONS

Conceptualization: D.T.V.D., M.B.; Investigation: D.T.V.D., J.F.C., M.H., M.W., O.S.; Visualization: M.B., D.T.V.D., J.F.C., M.H., M.W., O.S.; Writing – Original Draft: M.B.; Writing – Review and editing D.T.V.D., J.F.C., M.H., O.S., R.F.B, M.B.; Supervision: O.S., R.F.B, M.B.; Funding Acquisition: O.S., R.F.B, M.B.

## Abbreviations used in this paper

LD/LDs: lipid droplet(s)

vCLIP: vacuole and lipid droplet

NVJ: nucleus vacuole junction

kDa: kilodalton

Ldo16/45: lipid droplet organization protein of 16/45 kDa

LDAF1: lipid droplet assembly factor 1

Pdr16: pleiotropic drug resistance 16

Vac8: vacuole related protein 8

PITP: phosphatidylinositol transfer protein

