## Supplement for "A metabolically controlled contact site between lipid droplets and vacuoles"

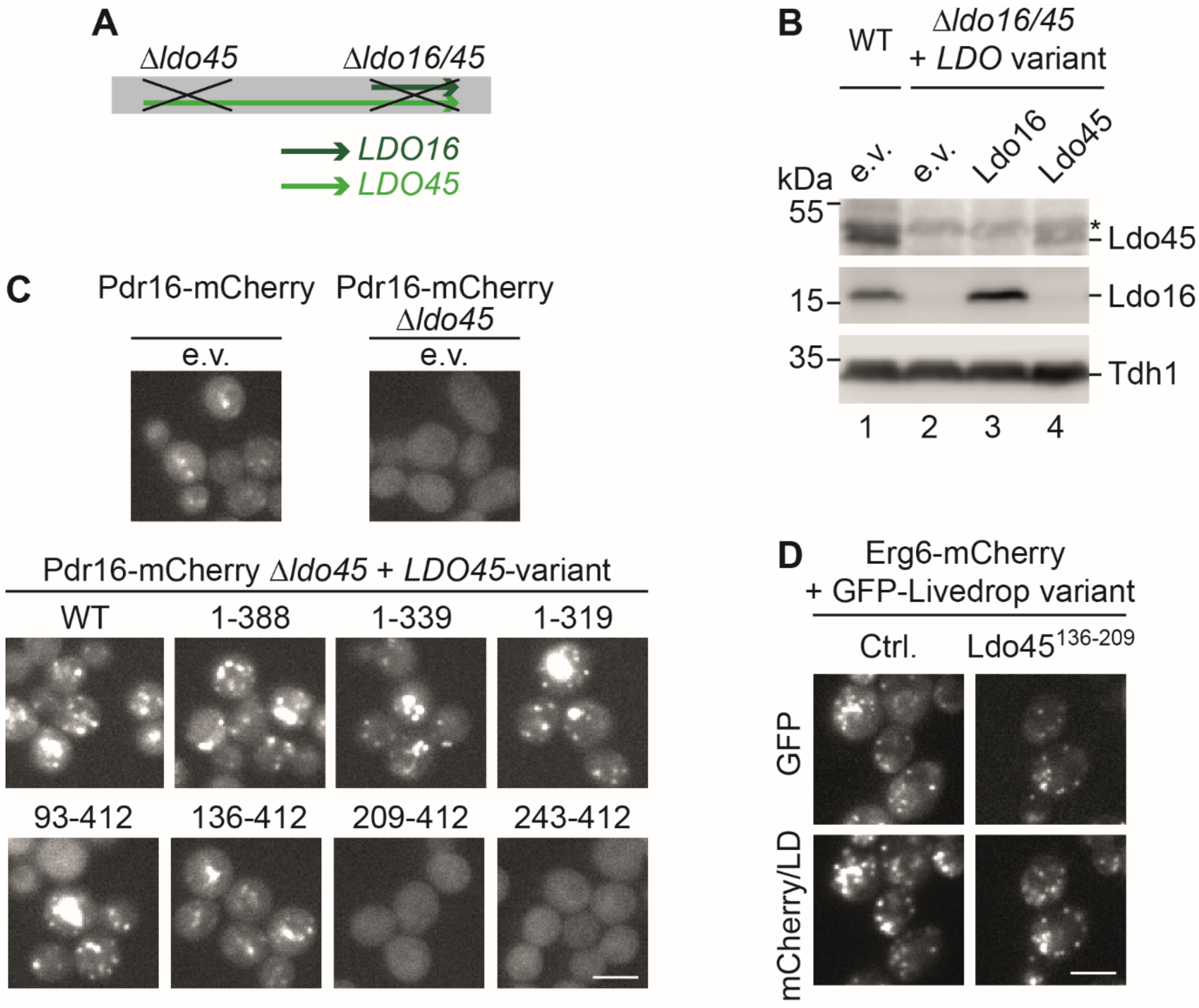


**Figure S1. Related to Figure 1. Structure-function analysis of Ldo45.**

(A) Schematic representation of the *LDO16/45* locus.

(B) Δ*ldo16/45* and wild-type (WT) control cells were transformed with centromeric plasmids for expression of either Ldo16 or Ldo45 under control of their own promoters or empty vectors (e.v.) and grown to logarithmic growth phase on synthetic medium supplemented with 2% glucose. Proteins were extracted and analyzed by SDS-PAGE and Western blotting. Asterisk: non-specific band.

(C) Indicated Ldo45 variants were expressed under control of the native *LDO45* promoter from centromeric plasmids in Pdr16-mCherry cells carrying a genomic *LDO45* deletion. Cells were cultured to logarithmic growth phase in synthetic medium containing 2% glucose and analyzed by fluorescence microscopy. e.v., empty vector. Scale bar, 5 µm.

(D) Erg6-mCherry cells expressing control (Ctrl.) LiveDrop or a LiveDrop variant fused to an Ldo45 domain (amino acids 136-209) from a centromeric plasmid under control of the *TEF2* promoter were cultured to logarithmic growth phase. Scale bar, 5 µm.

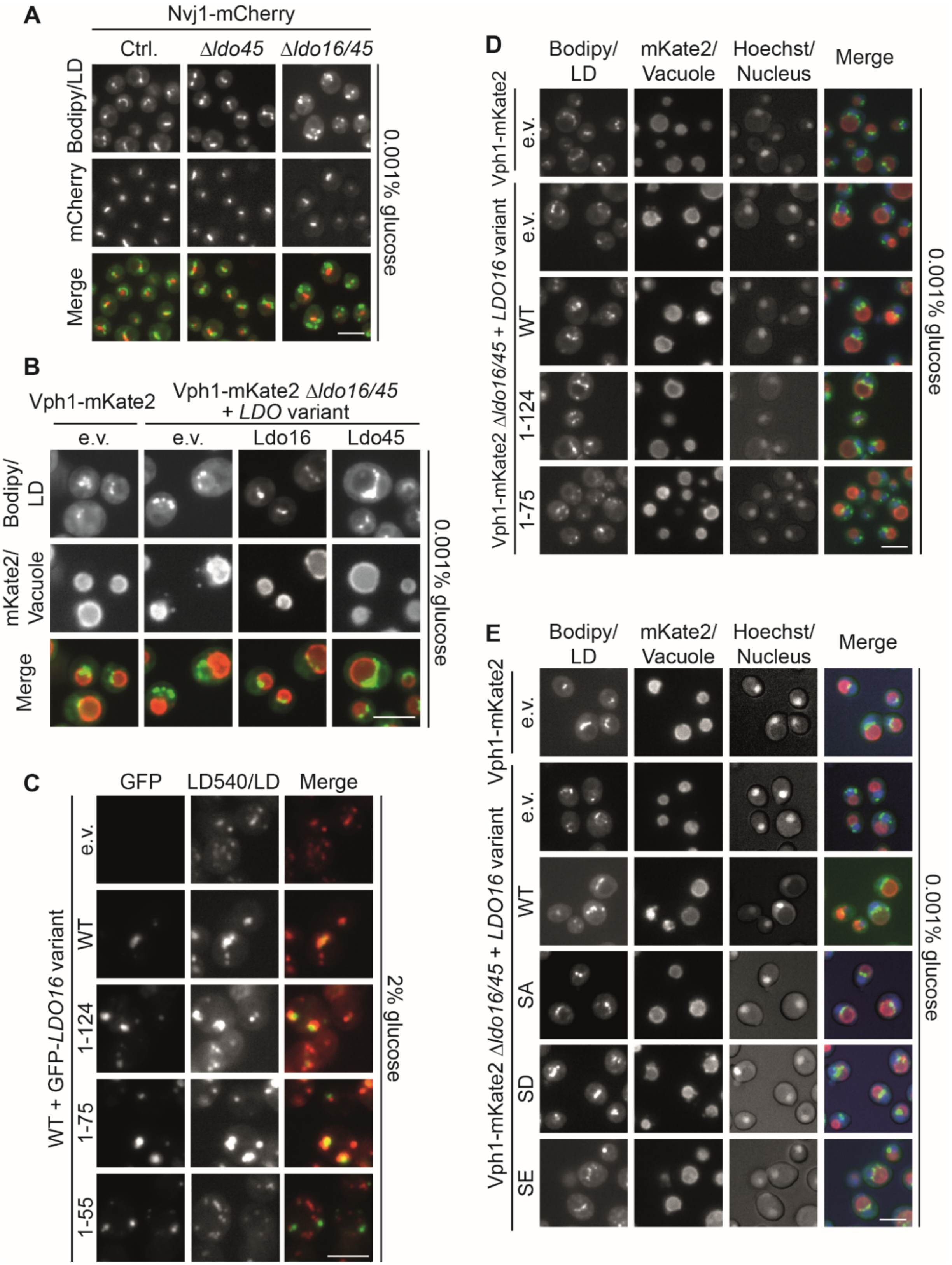

A
Nvj1-mCherry
Ctrl.
Δldo45
Δldo16/45
Bodipy/LD
mCherry
Merge
0.001% glucose
B
Vph1-mKate2
Vph1-mKate2 Δldo16/45 + LDO variant
e.v.
e.v.
Ldo16
Ldo45
Bodipy/ LD
mKate2/ Vacuole
Merge
0.001% glucose
C
GFP
LD540/LD
Merge
WT + GFP-LDO16 variant
e.v.
WT
1-124
1-75
1-55
2% glucose
D
Bodipy/ LD
mKate2/ Vacuole
Hoechst/ Nucleus
Merge
Vph1-mKate2
Vph1-mKate2 Δldo16/45 + LDO16 variant
e.v.
e.v.
WT
1-124
1-75
0.001% glucose
E
Bodipy/ LD
mKate2/ Vacuole
Hoechst/ Nucleus
Merge
Vph1-mKate2
Vph1-mKate2 Δldo16/45 + LDO16 variant
e.v.
e.v.
WT
SA
SD
SE
0.001% glucose

**Figure S2. Related to Figure 2. Structure-function analysis of Ldo16.**

(A) Control (Ctrl.), Δ*ldo45,* and Δ*ldo16/45* cells expressing the NVJ marker Nvj1-mCherry were grown overnight in synthetic medium containing 2% glucose, followed by a 4 hour incubation in 0.001% glucose and LDs were stained with BODIPY493/503 (Bodipy). LDs accumulated at the NVJ in the presence of LDO proteins, an effect that was abolished in Δ*ldo16/45* cells. Scale bar, 5 µm.

(B) Vph1-mKate2 Δ*ldo16/45* cells were transformed with centromeric plasmids for expression of Ldo16 or Ldo45 under control of their own promoters or with an empty vector (e.v.) and analyzed as described in (A). Scale bar, 5 µm.

(C) Indicated GFP-Ldo16 variants were expressed under control of a *TEF2* promoter from centromeric plasmids in wild-type (WT) cells. Cells were cultured to logarithmic growth phase in synthetic medium containing 2% glucose, stained with the neutral lipid dye LD540, and analyzed by fluorescence microcopy. All Ldo16 variants colocalized with LDs labeled by LD540 apart from the shortest variant, Ldo16$^{1\text{-}55}$, which formed foci of unknow identity distinct from LDs. Scale bar, 5 µm.

(D) Vph1-mKate2 Δ*ldo16/45* cells expressing full length Ldo16 (WT) and indicated truncated variants from centromeric plasmids under control of the *LDO16* promoter were analyzed as described in (A). Scale bar, 5 µm.

(E) Vph1-mKate2 Δ*ldo16/45* cells expressing native Ldo16 (WT), non-phosphorylatable Ldo16$^{S102A}$ (SA) or phosphomimetic Ldo16$^{S102D}$ (SD) and Ldo16$^{S102E}$ (SE) variants from centromeric plasmids under control of the *LDO16* promoter were analyzed as described in (A). Scale bar, 5 µm.

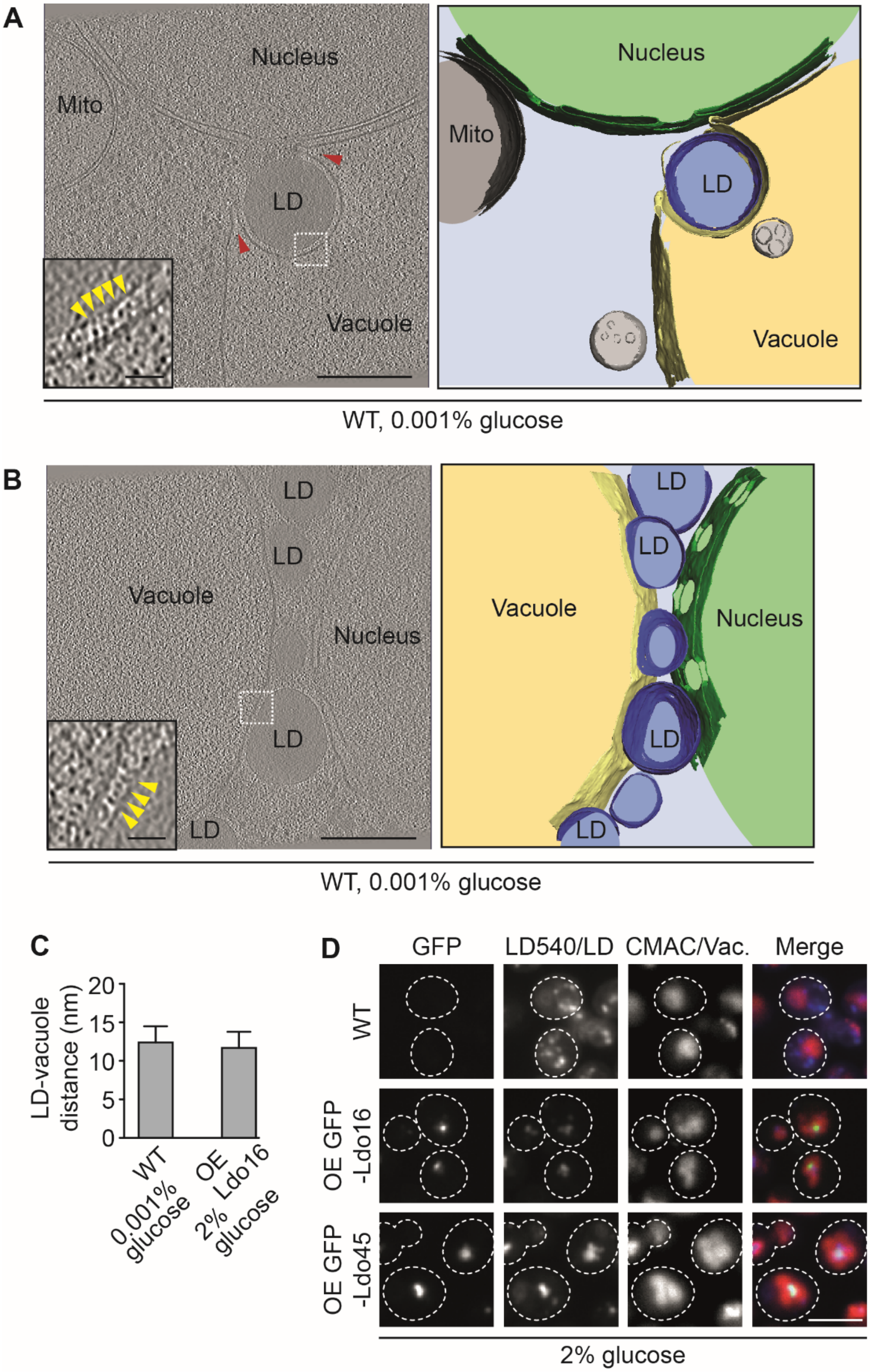
A
Nucleus
Mito
LD
Vacuole
WT, 0.001% glucose
B
LD
Vacuole
Nucleus
WT, 0.001% glucose
C
LD-vacuole distance (nm)
20
15
10
5
0
WT 0.001% glucose
OE Ldo16 2% glucose
D
GFP
LD540/LD
CMAC/Vac.
Merge
WT
OE GFP -Ldo16
OE GFP -Ldo45
2% glucose

**Figure S3. Related to Figures 2 and 3. LDs and vacuoles are tightly linked by contact sites.**

(A-B) Tomograms of vacuole-LD contact sites in wild-type (WT) cells cultured overnight in synthetic medium containing 2% glucose, followed by an incubation in 0.001% glucose for an additional 4 hours. Scale bar, 200 nm. Red arrowheads indicate vacuolar membrane deformations. Inset shows electron-dense objects at the LD-vacuole interface that may represent molecular tethers (indicated by yellow arrowheads). Scale bar, 25 nm. 3D segmentation depicted on the right. Mito, mitochondrion.

(C) The distance between LDs and vacuoles at contact sites was measured in tomograms of wild-type (WT) cells cultured overnight in synthetic medium containing 2% glucose, followed by a 4 hour incubation in 0.001% glucose (left). Cells overexpressing Ldo16 and grown to logarithmic growth phase in synthetic medium containing 2% glucose were analyzed in the same way (right). n=100. Error bars indicate SD values.

(D) GFP-Ldo16 and GFP-Ldo45 were overexpressed from a *TEF2* promoter. LDs (labeled by LD540, displayed in blue) colocalized with vacuoles (stained by CMAC, displayed in red) upon LDO overexpression, but not in wild-type (WT) controls. Scale bar, 5 µm.

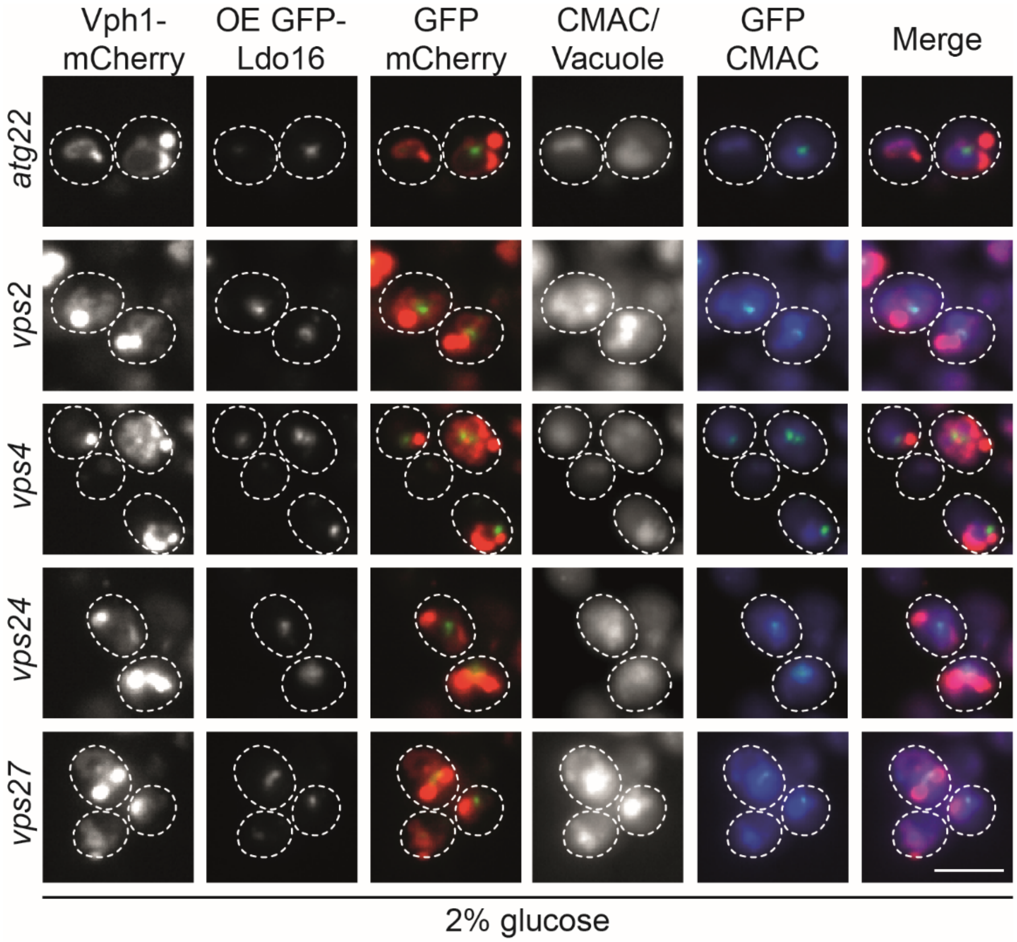


**Figure S4. Related to Figure 4. Identification of components affecting vCLIP formation.**

Mutants identified as hits belonging to the class “ambiguous” in the screen for genes affecting vCLIPs induced by overexpression of GFP-Ldo16 described in Figures 4A and B were cultured to logarithmic growth phase in synthetic medium containing 2% glucose, stained with the vacuolar lumen dyes CMAC and analyzed by fluorescence microscopy. CMAC staining revealed that for the indicated mutants, loss of colocalization of GFP-Ldo16 and the vacuolar membrane marker Vph1-mCherry was not due to a vCLIP defect, but caused by a mislocalization of Vph1-mCherry to non-vacuole structures. Scale bar, 5 µm.

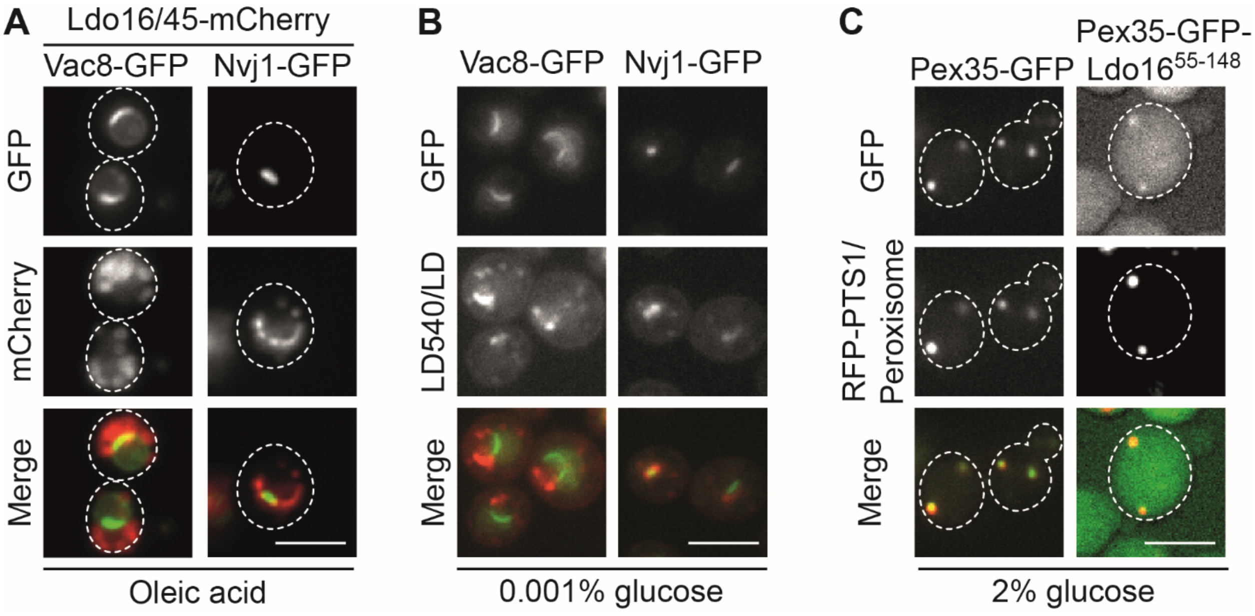


**Figure S5. Related to Figure 5. Vac8 is required for vCLIP formation.**

(A) Ldo16/45-mCherry Vac8-GFP and Ldo16/45-mCherry Nvj1-GFP cells were grown in synthetic medium supplemented with 2% glucose overnight, and further cultured in glucose-free medium supplemented with 0.2% oleic acid for an additional 24 hours. Scale bar, 5 µm.

(B) Vac8-GFP and Nvj1-GFP cells were cultured overnight in synthetic medium containing 2% glucose, followed by a 4 hour incubation in 0.001% glucose, LD540 staining, and fluorescence microscopy. Scale bar, 5 µm.

(C) Wild-type (WT) cells were transformed with plasmids encoding Pex35-GFP or Pex35-GFP-Ldo16$^{55-148}$ and an RFP-PTS1 plasmid encoding a fluorescent peroxisome marker, grown to logarithmic growth phase in synthetic medium containing 2% glucose, and analyzed by fluorescence microscopy. Scale bar, 5 µm.

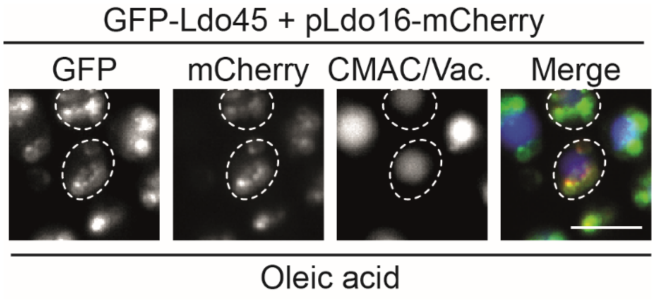


**Figure S6. Related to Figure 6. Ldo16 and Ldo45 localize simultaneously to vCLIP.**

Cells expressing GFP-Ldo45 under control of the *LDO45* promoter from the genome were transformed with a centromeric plasmid for expression of Ldo16-mCherry under control of the *LDO16* promoter, cultured in synthetic medium containing 2% glucose overnight, and then grown in glucose-free medium supplemented with 0.2% oleic acid for an additional 24 hours, stained with the vacuole marker CMAC, and analyzed by fluorescence microscopy. Both Ldo16 and Ldo45 formed foci at vCLIPs. Scale bar, 5 μm.

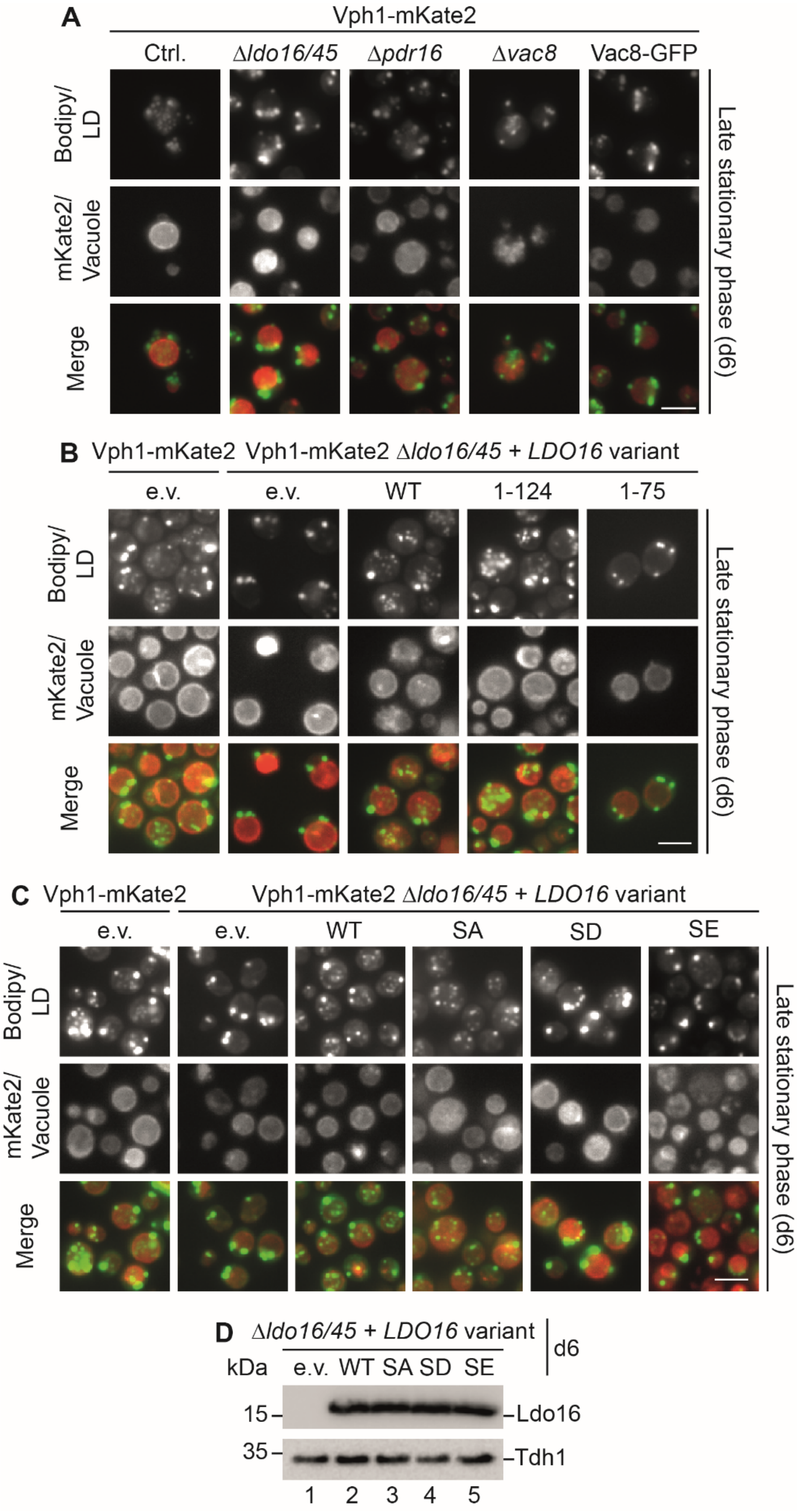
A
Vph1-mKate2
Ctrl.
Δldo16/45
Δpdr16
Δvac8
Vac8-GFP
Bodipy/ LD
mKate2/ Vacuole
Merge
Late stationary phase (d6)
B
Vph1-mKate2
Vph1-mKate2 Δldo16/45 + LDO16 variant
e.v.
e.v.
WT
1-124
1-75
Bodipy/ LD
mKate2/ Vacuole
Merge
Late stationary phase (d6)
C
Vph1-mKate2
Vph1-mKate2 Δldo16/45 + LDO16 variant
e.v.
e.v.
WT
SA
SD
SE
Bodipy/ LD
mKate2/ Vacuole
Merge
Late stationary phase (d6)
D
Δldo16/45 + LDO16 variant
d6
kDa
e.v. WT SA SD SE
15
Ldo16
35
Tdh1
1 2 3 4 5

**Figure S7. Related to Figure 7. Efficient lipophagy depends on the C-terminal domain of Ldo16 and correlates with dephosphorylation of Ldo16 S102.**

(A) Control (Ctrl.), $\Delta$*ldo16/45*, $\Delta$*pdr16*, $\Delta$*vac8*, and Vac8-GFP cells expressing the vacuolar membrane marker Vph1-mKate2 were cultured overnight in synthetic medium containing 2% glucose, diluted in fresh medium, and grown for another 6 days to induce lipophagy. LDs were visualized by BODIPY493/503 (Bodipy). e.v., empty vector. Scale bar, 5 µm.

(B) Vph1-mKate2 $\Delta$*ldo16/45* cells expressing indicated Ldo16 variants (full length/WT as well as truncated variants) from centromeric plasmids under control of the *LDO16* promoter were analyzed as described in (A). Scale bar, 5 µm.

(C) Vph1-mKate2 $\Delta$*ldo16/45* cells expressing indicated Ldo16 variants (native/WT as well as the non-phosphorylatable variant Ldo16$^{S102A}$ (SA) and the phosphomimetic variants Ldo16$^{S102D}$ (SD) and Ldo16$^{S102E}$ (SE)) from centromeric plasmids under control of the *LDO16* promoter were analyzed as described in (A). Scale bar, 5 µm.

(D) $\Delta$*ldo16/45* cells were transformed with centromeric plasmids for expression of Ldo16 variants (native/WT as well as the non-phosphorylatable variant Ldo16$^{S102A}$ (SA) and the phosphomimetic variants Ldo16$^{S102D}$ (SD) and Ldo16$^{S102E}$ (SE)) under control of the *LDO16* promoter. Proteins were extracted on day 6 (d6) and analyzed by SDS-PAGE and Western blotting.

### MATERIALS AND METHODS

*Yeast strains and growth condition*

The yeast strains used in this study are derivatives of the *Saccharomyces cerevisiae* wild-type strain BY4741 (*MATa his3Δ1 leu2Δ0 met15Δ0 ura3Δ0*) (Brachmann et al., 1998) and a related wild-type carrying features required for automated mutant generation (*MATα his3Δ1 leu2Δ0 met15Δ0 ura3Δ0 can1Δ::STE2pr-spHIS5 lyp1Δ::STE3pr-LEU2*) (Breslow et al., 2008). Cells were genetically manipulated by PCR-based homologous recombination (Longtine et al., 1998). Primers for genetic modification and validation were designed using Primers-4-Yeast (Yofe and Schuldiner, 2014). Yeast cells were pre-cultured overnight in synthetic medium (0.67 % weight/volume yeast nitrogen base with ammonium sulfate, 2% weight/volume glucose, amino acid supplements: alanine, arginine, asparagine, aspartic acid, cysteine, glutamine, glutamic acid, glycine, inositol, isoleucine, lysin, methionine, para-aminobenzoic acid, phenylalanine, proline, serine, threonine, tyrosine, valine, histidine, leucine, uracil, tryptophan, adenine) in a shaking incubator at 30°C, 280 rpm. On the next day, they were typically diluted and grown until they reached the logarithmic phase. For analysis of cells in early stationary phase, cells were kept undiluted. For glucose restriction, overnight-cultured cells in 2% glucose were collected, washed two times with synthetic medium containing 0.001% glucose, resuspended in the same medium, and grown for a further 4 hours. For induction of lipophagy, cells were cultured overnight in synthetic medium containing 2% glucose, followed by a dilution in the same medium. Cells were analyzed after 1, 4, and 6 days of growth. For oleic acid treatment, overnight-cultured cells in 2% glucose were collected, washed two times in glucose-free medium containing 0.2% oleic acid and 0.1% Tween 80, and resuspended in the same medium. Subsequently, cells were grown for an additional 24 hours. For the inositol-depletion condition, cells were cultured overnight and diluted in inositol-free synthetic medium supplemented with 2% glucose.

*Generation of plasmids*

Plasmids were constructed by the Gibson Assembly cloning method. PCR fragments and a vector with appropriate overlapping ends were mixed with NEBuilder HiFi DNA Assembly Master Mix (NEB), followed by an incubation at 50°C for 15 minutes. Subsequently, ligated constructs were transformed into competent Escherichia coli cells, followed by DNA extraction and sequencing.

*Fluorescence microscopy*

Cells were transferred to a 384-well microscope plate (Brooks) with a glass bottom coated with Concanavalin A (Sigma-Aldrich). After 15 minutes, cells were washed with respective medium. For staining of organelles, respective dyes were added at this step. For LD staining, cells were incubated with BODIPY 493/503 (1 µM; Sigma-Aldrich) or LD540 (0.5 µg/mL) for 15 minutes or 30 minutes, respectively. For vacuole staining, cells were incubated with CMAC (0.1 mM; ThermoFisher) for 30 minutes. Alternatively, cells were incubated with FM4-64 (8 µM; ThermoFisher) for 30 minutes, followed by washing with respective medium and incubation for a further 30 minutes at 30°C. For nucleus staining, Hoechst (6 µg/mL; ThermoFisher) was added for 30 minutes. After indicated incubation time for each dye, cells were washed with medium without dye and ready for imaging. Cells were imaged with the Olympus screening station ScanR at room temperature. Images were taken with the Olympus ScanR Automated Image Acquisition Software using the Olympus IX83 inverted fluorescence microscope with the Lumencor SpectraX LED light source and a 40x or 60x air objective. Images were processed using ImageJ.

*Automated library preparation and high content imaging*

Two query strains (Vph1-mCherry pTEF2-GFP-Ldo16 and Vph1-mKate2) were created in a background carrying genetic features required for synthetic genetic arrays and crossed with a genome wide deletion (Giaever et al., 2002) and hypomorphic allele (Breslow et al., 2008) mutant collection using an automated mating protocol (Cohen and Schuldiner, 2011; Tong and Boone, 2006). Handling of the systematic mutant collections was performed using the RoToR benchtop colony array instrument (Singer instruments). Strains were mated and, after selection of diploids, sporulation was induced by transfer to nitrogen starvation medium. For haploid selection, strains were replicated on plates containing 50 mg/L Canavanine (Sigma-Aldrich) and 50 mg/L Thialysine (Sigma-Aldrich). For the final selection step, a combination of all selections of the desired mutations was used. Before imaging, strains were grown in 384 well polysterene plates at the indicated growth conditions, stained with Bodipy if required, and then moved to 384 well glass bottom plates (Brooks) using the Bench Smart 96TM liquid handler (Mettler Toledo). Automated imaging of mutant collection plates was performed using the ScanR System (Olympus).

*Whole cell extraction, SDS-PAGE and Western blotting*

Cells were grown to logarithmic growth phase and harvested by centrifugation. Subsequently, proteins were extracted via alkaline lysis (Kushnirov, 2000) and subjected to SDS-PAGE and Western blotting. Membranes were analyzed by enhanced chemiluminescence (ECL) using the Azure imaging system.

*Phos-tag PAGE*

Samples were prepared by an adapted alkaline lysis protocol (Kushnirov, 2000) as described previously (Schmidt et al., 2019). Cells were harvested by centrifugation, and washed once with ice-cold phosphatase inhibitor mix (10 mM NaF, 10 mM β-glycerol phosphate, PhosSTOP; Roche, 1 tablet per 100 ml). After centrifugation (3 minutes, 15,000 × g, 4°C) cells were resuspended in 0.1 M NaOH with the same phosphatase inhibitors and incubated at room temperature for 5 minutes. Finally, cells were recovered by centrifugation and resuspended in Lämmli sample buffer (60mM Tris/HCl pH 6.8, 2% SDS, 10% glycerol, 1% beta-mercaptoethanol, 0.01% bromo phenol blue), denatured (95°C, 15 minutes), and cell debris was removed by centrifugation (3 minutes, 15,000 × g, 4°C). The cleared lysate was loaded onto 12.5% Phos-tag SDS PAGE gels prepared according to the manufacturer's specifications. 50 μM Phos-tag acrylamide (Wako); 100 μm

$MnCl_2$ were used for detection of Ldo16/45-FLAG and 35 µM Phostag/70 µM $MnCl_2$ for detection with the Ldo16/45 antibody. Gels were run in a standard Lämmli electrophoresis buffer at 200V, 40 mA for 55 minutes, afterwards rinsed in Western blot transfer buffer (25 mM Tris, 192 mM glycine, 20% ethanol, 0.1% SDS) with 10 mM EDTA for 20 minutes, and equilibrated in transfer buffer (5 minutes), followed by wet electroblotting to PVDF membranes (80V constant, 2 hrs). Membranes were analyzed by enhanced chemiluminescence (ECL) (Advansta Western bright) using the Vilber fusion FX7 edge imaging system.

*GFP immunoprecipitation*

Equal amounts of cells expressing GFP-tagged proteins and untagged controls were harvested via centrifugation. Samples were resuspended with 500 µl glass beads (Sigma-Aldrich) and 500 µl GFP pulldown buffer (150 mM KOAc, 20 mM HEPES pH 7.4, 5% glycerol, complete protease inhibitor cocktail (Roche), 1% Octyl-beta-Glucoside (ThermoFisher)). Cells were lysed using a shaker (IKA VIBRAX). Cleared lysates were collected and 50 µl of each samples was reserved as “input”. The remaining cell lysates were incubated with pre-equilibrated GFP-Trap agarose beads (Chromotek) for 60 minutes at 4°C rotating. Subsequently, beads were washed twice GFP pulldown buffer and four times in the same buffer without detergent. After washing, beads were incubated with 50 µl HU buffer at 65°C for 15 minutes for elution. Input samples were treated in the same way with 50 µl HU buffer. Eluate and input samples were subjected to SDS-PAGE and Western blotting.

*Cryo-electron tomography*

Cryo-EM grids (R1.2/1.3, Cu 200 mesh grid, Quantifoil microtools) were glow discharged in a plasma cleaner (PDC-32G-2, Harrick Plasma) with their carbon side facing up for 30 s on medium setting. The grids were mounted on a Vitrobot Mark IV (Thermo Fisher Scientific) and a 3.5 µl drop of yeast culture at 0.8 $OD_{600}$ was deposited on their carbon side. Excess liquid was removed by back-blotting with filter paper (Whattman 597). The grids were quickly plunged into a liquid ethane/propane mixture at liquid nitrogen temperature. The grids were stored in grid boxes and submerged in liquid nitrogen until use.

Vitrified grids were mounted into Autogrid carriers (Thermo Fisher Scientific) and secured in place by a copper ring. They were subsequently loaded in a 35° tilt shuttle (Thermo Fisher Scientific) and inserted in an Aquilos 2 Cryo focused ion beam/scanning electron microscope (Thermo Fisher Scientific) where they were kept below -180°C by the microscope's cryo-stage. Single-layer clusters of cells were targeted for milling for their higher chance of good vitrification. A layer of organometallic platinum was deposited on top of the grid with the microscope's gas injection system for 40 seconds to protect the peripheral areas of the milling target region from the ion beam. Milling was done at a sample tilt of 20°. The milling took place sequentially for each position, starting with the $Ga^{2+}$ ion beam at 30 kV and 300 pA beam current for rough milling and ending at 30 kV and 50 pA for fine milling. The target dimensions for the lamellae were 15 μm wide and 200 nm thick. SEM imaging at 3 kV and 13 pA was used to monitor the milling process.

The lamellae were inserted in a Krios G4 Cryo- transmission electron microscope (Thermo Fisher Scientific) with a 300kV field emission gun, Selectris energy filter, and a Falcon 4i direct electron detector camera. An overall map of the grid was acquired at very low magnification to identify the positions of the lamellae. Afterwards, low magnification overviews of the lamellae were taken to identify regions of interest within them. Tilt series were recorded using SerialEM software (Mastronarde, 2005) at higher magnification (42,000 ×, 2.935 Å pixel size; -5 to -6 μm defocus), typically from -46° to +64° at increments of 3° using the dose-symmetric acquisition scheme (Hagen et al., 2017). The target total dose per tomogram was around 100 e-/A$^2$, which translated to a ~2.8 s exposure per tilt image. The camera was operated in dose-fractionation mode and between 700 to 900 EER frames were generated for each tilt image.

The raw EER frames were processed by SNARTomo (https://github.com/rubenlab/snartomo), which uses Motioncor2 (Zheng et al., 2017) to align the frames. The resulting new tilt series was later reconstructed using IMOD (Kremer et al., 1996) with patch-tracking as tilt image alignment method and weighted back-projection for the final tomogram reconstruction. Tomograms were binned twice to a final pixel size of 1.17 nm. Where specified, tomograms were denoised using Topaz 3D (Bepler et al., 2020).

The membranes in selected tomograms were segmented to obtain 3D renderings. The software TomoSegMemTV (Martinez-Sanchez et al., 2014) was used for the initial automatic tracing of the membranes, and Amira (Thermo Fisher Scientific) for their refinement and coloring.

*Quantifications, statistical analysis*

In Figures 2B and D, cells were visually classified based on the spatial relationship between Bodipy-labeled LDs and vacuoles marked by Vph1-mKate2 into two groups: cells with LDs accumulating on the vacuolar membrane, and cells with LDs far from vacuolar membranes. n=100 cells. In Figure 6C, protein levels of Ldo16-FLAG and Ldo45-FLAG normalized to the Pgk1 loading control were represented as fold change from logarithmic growth phase. n=3. Error bars indicate SD values. In Figures 7B, C, D and E, the number of Bodipy-stained LDs located inside the vacuole (marked by Vph1-mKate2) was counted and cells were classified into three groups: cells with 0 LDs/vacuole, cells with 1-3 LDs/vacuole, and cells with >3 LDs/ vacuole. n=100 cells (in Figures 7C, D, E) or 50 cells (in Figure 7B). In Figure 7F, the ratio between phosphorylated Ldo16-FLAG and dephosphorylated Ldo16-FLAG determined. n=3. Error bars indicate SD values.

### List of yeast strains used in this study

| *Name and genotype* | *Source* | *Identifier* |
|---|---|---|
| WT - MATα *his3Δ1 leu2Δ0 met15Δ0 ura3Δ0 can1Δ::STE2pr-spHIS5 lyp1Δ::STE3pr-LEU2* | Breslow et al., 2008 | yMB3 |
| *△ldo16/45* - MATα *his3Δ1 leu2Δ0 met15Δ0 ura3Δ0 can1Δ::STE2pr-spHIS5 lyp1Δ::STE3pr-LEU2 △ldo16/45::NAT* | Eisenberg-Bord et al., 2018 | yMB114 |
| *△sei1*- MATα *his3Δ1 leu2Δ0 met15Δ0 ura3Δ0 can1Δ::STE2pr-spHIS5 lyp1Δ::STE3pr-LEU2 △sei1::Hygro* | This study | yMB244 |
| *△ldo45* - MATα *his3Δ1 leu2Δ0 met15Δ0 ura3Δ0 can1Δ::STE2pr-spHIS5 lyp1Δ::STE3pr-LEU2 △ldo45::NAT* | Eisenberg-Bord et al., 2018 | yMB97 |
| *△ldb16* - MATα *his3Δ1 leu2Δ0 met15Δ0 ura3Δ0 can1Δ::STE2pr-spHIS5 lyp1Δ::STE3pr-LEU2 △ldb16::Hygro* | This study | yMB250 |
| *△vac8* - MATα *his3Δ1 leu2Δ0 met15Δ0 ura3Δ0 can1Δ::STE2pr-spHIS5 lyp1Δ::STE3pr-LEU2 △vac8::NAT* | This study | yMB883 |
| *△pdr16* - MATα *his3Δ1 leu2Δ0 met15Δ0 ura3Δ0 can1Δ::STE2pr-spHIS5 lyp1Δ::STE3pr-LEU2 △pdr16::G418* | This study | yMB107 |
| Vac8-GFP - MATα *his3Δ1 leu2Δ0 met15Δ0 ura3Δ0 can1Δ::STE2pr-spHIS5 lyp1Δ::STE3pr-LEU2 Vac8-GFP::G418* | This study | yMB885 |
| Pdr16-GFP - MATα *his3Δ1 leu2Δ0 met15Δ0 ura3Δ0 can1Δ::STE2pr-spHIS5 lyp1Δ::STE3pr-LEU2 Pdr16-GFP::G418* | Eisenberg-Bord et al., 2018 | yMB18 |
| Pdr16-mCherry - MATα *his3Δ1 leu2Δ0 met15Δ0 ura3Δ0 can1Δ::STE2pr-spHIS5 lyp1Δ::STE3pr-LEU2 Pdr16-mCherry::NAT* | Eisenberg-Bord et al., 2018 | yMB9 |
| Pdr16-mCherry *△ldo45* - MATα *his3Δ1 leu2Δ0 met15Δ0 ura3Δ0 can1Δ::STE2pr-spHIS5 lyp1Δ::STE3pr-LEU2 Pdr16-mCherry::NAT △ldo45::G418* | Eisenberg-Bord et al., 2018 | yMB80 |
| Ldo16-GFP Erg6-mCherry - MATα *his3Δ1 leu2Δ0 met15Δ0 ura3Δ0 can1Δ::STE2pr-spHIS5 lyp1Δ::STE3pr-LEU2 Ldo16-GFP::HIS Erg6-mCherry::NAT* | This study | yMB120 |
| Ldo16-GFP Erg6-mCherry *△vac8* - MATα *his3Δ1 leu2Δ0 met15Δ0 ura3Δ0 can1Δ::STE2pr-spHIS5 lyp1Δ::STE3pr-LEU2 Ldo16-GFP::HIS Erg6-mCherry::NAT △vac8::G418* | This study | yMB884 |
| Ldo16-GFP Erg6-mCherry *△sei1* - MATα *his3Δ1 leu2Δ0 met15Δ0 ura3Δ0 can1Δ::STE2pr-spHIS5 lyp1Δ::STE3pr-LEU2 Ldo16-GFP::HIS Erg6-mCherry::NAT △sei1::Hygro* | This study | yMB915 |
| Pdr16-GFP Erg6-mCherry - MATα *his3Δ1 leu2Δ0 met15Δ0 ura3Δ0 can1Δ::STE2pr-spHIS5 lyp1Δ::STE3pr-LEU2 Pdr16-GFP::G418 Erg6-mCherry::NAT* | Eisenberg-Bord et al., 2018 | yMB77 |

| | | |
|---|---|---|
| Pdr16-GFP Erg6-mCherry △*ldo45*- MATα *his3Δ1 leu2Δ0 met15Δ0 ura3Δ0 can1Δ::STE2pr-spHIS5 lyp1Δ::STE3pr-LEU2 Pdr16-GFP::G418 Erg6-mCherry::NAT △ldo45::Hygro* | This study | yMB916 |
| Pdr16-GFP Ldo16-mCherry - MATα *his3Δ1 leu2Δ0 met15Δ0 ura3Δ0 can1Δ::STE2pr-spHIS5 lyp1Δ::STE3pr-LEU2 Pdr16-GFP:G418 Ldo16-mCherry::NAT* | This study | yMB914 |
| Pdr16-GFP Ldo16-mCherry △*ldo45* - MATα *his3Δ1 leu2Δ0 met15Δ0 ura3Δ0 can1Δ::STE2pr-spHIS5 lyp1Δ::STE3pr-LEU2 Pdr16-GFP:G418 Ldo16-mCherry::NAT △ldo45::Hygro* | This study | yMB938 |
| TEF2p-GFP-Ldo16 - MATα *his3Δ1 leu2Δ0 met15Δ0 ura3Δ0 can1Δ::STE2pr-spHIS5 lyp1Δ::STE3pr-LEU2 TEF2p-GFP-Ldo16::NAT* | Eisenberg-Bord et al., 2018 | yMB58 |
| TEF2p-GFP-Ldo45 - MATα *his3Δ1 leu2Δ0 met15Δ0 ura3Δ0 can1Δ::STE2pr-spHIS5 lyp1Δ::STE3pr-LEU2 TEF2p-GFP-Ldo45::NAT* | Eisenberg-Bord et al., 2018 | yMB57 |
| TEF2p-Vac8 - MATα *his3Δ1 leu2Δ0 met15Δ0 ura3Δ0 can1Δ::STE2pr-spHIS5 lyp1Δ::STE3pr-LEU2 TEF2p-Vac8::NAT* | This study | yMB924 |
| Vph1-mKate2 - MATα *his3Δ1 leu2Δ0 met15Δ0 ura3Δ0 can1Δ::STE2pr-spHIS5 lyp1Δ::STE3pr-LEU2 Vph1-mKate2::G418* | Eising et al., 2022 | yMB352 |
| Vph1-mCherry TEF2p-GFP-Ldo16 - MATα *his3Δ1 leu2Δ0 met15Δ0 ura3Δ0 can1Δ::STE2pr-spHIS5 lyp1Δ::STE3pr-LEU2 Vph1-mCherry::HIS TEF2p-GFP-Ldo16::NAT* | This study | yMB888 |
| Vph1-mKate2 △*ldo16/45* - MATα *his3Δ1 leu2Δ0 met15Δ0 ura3Δ0 can1Δ::STE2pr-spHIS5 lyp1Δ::STE3pr-LEU2 Vph1-mKate2::G418 △ldo16/45::Hygro* | This study | yMB612 |
| Vph1-mKate2 △*vac8* - MATα *his3Δ1 leu2Δ0 met15Δ0 ura3Δ0 can1Δ::STE2pr-spHIS5 lyp1Δ::STE3pr-LEU2 Vph1-mKate2::Hygro △vac8::NAT* | This study | yMB935 |
| Vph1-mKate2 △*pdr16* - MATα *his3Δ1 leu2Δ0 met15Δ0 ura3Δ0 can1Δ::STE2pr-spHIS5 lyp1Δ::STE3pr-LEU2 Vph1-mKate2::Hygro △pdr16::G418* | This study | yMB941 |
| Vph1-mKate2 Vac8-GFP - MATα *his3Δ1 leu2Δ0 met15Δ0 ura3Δ0 can1Δ::STE2pr-spHIS5 lyp1Δ::STE3pr-LEU2 Vph1-mKate2::Hygro Vac8-GFP::G418* | This study | yMB933 |
| TEF2p-Ldo16 Faa4-GFP - MATα *his3Δ1 leu2Δ0 met15Δ0 ura3Δ0 can1Δ::STE2pr-spHIS5 lyp1Δ::STE3pr-LEU2 TEF2p-Ldo16::NAT Faa4-GFP::HIS* | This study | yMB224 |
| TEF2p-Ldo16 Faa4-GFP △*vac8* - MATα *his3Δ1 leu2Δ0 met15Δ0 ura3Δ0 can1Δ::STE2pr-spHIS5 lyp1Δ::STE3pr-LEU2 TEF2p-Ldo16::NAT Faa4-GFP::HIS △vac8::G418* | This study | yMB873 |
| GFP-Ldo45 - MATα *his3∆1 leu2∆0 met15∆0 ura3∆0 lys+ can1∆::GAL1pr-SceI::STE2pr-SpHIS5 lyp1∆::STE3pr-LEU2 YMR147Wp-sfGFP-YMR147W* | Weill et al., 2016 | yMB82 |

| | | |
|---|---|---|
| Nvj1-mCherry△*ldo16/45* - MATα *his3Δ1 leu2Δ0 met15Δ0 ura3Δ0 can1Δ::STE2pr-spHIS5 lyp1Δ::STE3pr-LEU2 Nvj1-mCherry::Hygro △ldo16/45::NAT* | Eisenberg-Bord et al., 2018 | yMB136 |
| Nvj1-mCherry △*ldo45* - MATα *his3Δ1 leu2Δ0 met15Δ0 ura3Δ0 can1Δ::STE2pr-spHIS5 lyp1Δ::STE3pr-LEU2 Nvj1-mCherry::Hygro △ldo45::NAT* | Eisenberg-Bord et al., 2018 | yMB102 |
| Nvj1-mCherry - MATα *his3Δ1 leu2Δ0 met15Δ0 ura3Δ0 can1Δ::STE2pr-spHIS5 lyp1Δ::STE3pr-LEU2 Nvj1-mCherry::NAT* | This study | yMB105 |
| Zrc1-VN Faa4-VC - MATα *his3Δ1 leu2Δ0 met15Δ0 ura3Δ0 can1Δ::STE2pr-spHIS5 lyp1Δ::STE3pr-LEU2 Zrc1-VN::HIS Faa4-VC::Hygro* | This study | yMB30 |
| Zrc1-VN Faa4-VC Ldo16-mCherry - MATα *his3Δ1 leu2Δ0 met15Δ0 ura3Δ0 can1Δ::STE2pr-spHIS5 lyp1Δ::STE3pr-LEU2 Zrc1-VN::HIS Faa4-VC::Hygro* Ldo16-mCherry::NAT | This study | yMB905 |
| Zrc1-VN Faa4-VC Pdr16-mCherry - MATα *his3Δ1 leu2Δ0 met15Δ0 ura3Δ0 can1Δ::STE2pr-spHIS5 lyp1Δ::STE3pr-LEU2 Zrc1-VN::HIS Faa4-VC::Hygro Pdr16-mCherry::NAT* | This study | yMB47 |
| Vac8-GFP Ldo16-mCherry - MATα *his3Δ1 leu2Δ0 met15Δ0 ura3Δ0 can1Δ::STE2pr-spHIS5 lyp1Δ::STE3pr-LEU2 Vac8-GFP::G418 Ldo16-mCherry::NAT* | This study | yMB904 |
| Nvj1-GFP Ldo16-mCherry - MATα *his3Δ1 leu2Δ0 met15Δ0 ura3Δ0 can1Δ::STE2pr-spHIS5 lyp1Δ::STE3pr-LEU2 Nvj1-GFP::G418 Ldo16-mCherry::HIS* | This study | yMB920 |
| Ldo16/45-FLAG - MATα *his3Δ1 leu2Δ0 met15Δ0 ura3Δ0 can1Δ::STE2pr-spHIS5 lyp1Δ::STE3pr-LEU2 Ldo16/45-FLAG::HIS* | This study | yMB881 |
| Deletion library | Giaever et al., 2002 | N/A |
| Decreased abundance by mRNA perturbation (DAmP) library | Breslow et al., 2008 | N/A |

### List of plasmids used in this study

| *Name* | *Source* | *Identifier* |
|---|---|---|
| pRS313 | Sikorski and Hieter, 1989 | pMB106 |
| pRS313-TEF2p-GFP-LDO16 | Eisenberg-Bord et al., 2018 | pMB42 |
| pRS313-TEF2p-GFP-LDO45 | Eisenberg-Bord et al., 2018 | pMB96 |
| pRS313-LDO16p-LDO16 | This study | pMB139 |
| pRS313-LDO45p-LDO45 | This study | pMB97 |
| pRS313-TEF2p-GFP-LDO45$^{1-319}$ | This study | pMB119 |
| pRS313-TEF2p-GFP-LDO45$^{1-339}$ | This study | pMB120 |
| pRS313-TEF2p-GFP-LDO45$^{1-388}$ | This study | pMB121 |
| pRS313-TEF2p-GFP-LDO45$^{243-412}$ | This study | pMB115 |
| pRS313-TEF2p-GFP-LDO45$^{209-412}$ | This study | pMB116 |
| pRS313-TEF2p-GFP-LDO451$^{36-412}$ | This study | pMB117 |

| | | |
|---|---|---|
| pRS313-TEF2p-GFP-LDO45$^{93-412}$ | This study | pMB118 |
| pRS313-LDO45p -LDO45$^{1-319}$ | This study | pMB103 |
| pRS313-LDO45p-LDO45$^{1-339}$ | This study | pMB104 |
| pRS313-LDO45p-LDO45$^{1-388}$ | This study | pMB105 |
| pRS313-LDO45p-LDO45$^{243-412}$ | This study | pMB99 |
| pRS313-LDO45p-LDO45$^{209-412}$ | This study | pMB100 |
| pRS313-LDO45p-LDO451$^{36-412}$ | This study | pMB101 |
| pRS313-LDO45p-LDO45$^{93-412}$ | This study | pMB102 |
| pRS313-TEF2p-GFP-LDO16$^{1-124}$ | This study | pMB246 |
| pRS313-TEF2p-GFP-LDO16$^{1-75}$ | This study | pMB247 |
| pRS313-TEF2p-GFP-LDO16$^{1-55}$ | This study | pMB248 |
| pRS313-LDO16p-LDO16$^{1-124}$ | This study | pMB189 |
| pRS313-LDO16p-LDO16$^{1-75}$ | This study | pMB188 |
| pRS313-TEF2p-GFP-Livedrop | This study | pMB140 |
| pRS313-TEF2p-GFP-Ldo45$^{136-209}$-LiveDrop | This study | pMB146 |
| pRS313-TEF2p-Ldo45$^{136-209}$-LiveDrop | This study | pMB249 |
| pRS313-TEF2p-GFP-LiveDrop-Ldo16$^{55-148}$ | This study | pMB197 |
| pRS313-TEF2p-Pex35-GFP | This study | pMB250 |
| pRS313-TEF2p-Pex35-GFP- Ldo16$^{55-148}$ | This study | pMB251 |
| pRS313-LDO16p-LDO16$^{S102A}$ | This study | pMB212 |
| pRS313-LDO16p-LDO16$^{S102D}$ | This study | pMB213 |
| pRS313-LDO16p-LDO16$^{S102E}$ | This study | pMB214 |
| pRS313-LDO16p-LDO16-mCherry | This study | pMB211 |
| pFA6a-KanMX6 | Longtine et al., 1998 | pMB5 |
| pFA6a-NAT-MX6 | Goldstein et al., 1999 | pMB6 |
| pFA6-Hygro | Goldstein et al., 1999 | pMB10 |
| pFa6a-cherry-NAT | Tomer Ravid | pMB15 |
| p4636 RFP-PTS1 | Jeffrey Gerst | pMB39 |
| pFA6a-GFP(S65T)-KanMX6 | Longtine et al., 1998 | pMB4 |
| pBS34-mCherry-KAN | Naama Barkai | pMB13 |
| pYM25-mKate-HygR | Roland Wedlich-Söldner | pMB253 |
| pFA6-FLAG-HIS | Schmidt lab | pMB252 |
| pFA6a-VN-His3MX6 | Sung et al., 2007 | pMB20 |
| pFA6a-VC-Hygro | This study | pMB41 |
| GFP-LiveDrop | Wang et al., 2016 | pmMB9 |

### List of primers used in this study

| *Sequence* | *Source* | *Identifier* |
|---|---|---|
| GCGGCTAATCAATTCTACTC | This study | prMB68 |
| TGGACTTGTTATTCCGTGTTCCCTACTTTTTTTGATA ATGcggatccccgggttaattaa | This study | prMB69 |
| TGCTAAACTTGCGAAAAATGTTTTTTTATTGCCGAGG TTAgaattcgagctcgtttaaac | This study | prMB70 |
| ATACCCTTATTGGGCATTTC | This study | prMB71 |
| ATTACCCGCTTTATTAGCAG | This study | prMB72 |
| AAACCTACAATCCAATCACG | This study | prMB48 |

| | | |
|---|---|---|
| AAGGTTTCAAGAAAATAAGATAAAGTGAATAGGAAG GATGcggatccccgggttaattaa | This study | prMB49 |
| TAGGTTTTAAAATTATATAGCGAGAAGTACAATTCTA TCAgaattcgagctcgtttaaac | This study | prMB50 |
| CAATGTATCCCGTCCATTAC | This study | prMB51 |
| AAATGGTCAGTTTTGCTCTG | This study | prMB52 |
| TTAATTAACCCGGGGATCCG | This study | prMB23 |
| ATACCCTTATTGGGCATTTC | This study | prMB93 |
| GACTACTGCTAATAAAGCGGGTAATAAGTTCCAGCT CTCTcggatccccgggttaattaa | This study | prMB94 |
| TGCTAAACTTGCGAAAAATGTTTTTTTATTGCCGAGG TTAgaattcgagctcgtttaaac | This study | prMB95 |
| GACTACTGCTAATAAAGCGGGTAATAAGTTCCAGCT CTCTGGTCGACGGATCCCCGGG | This study | prMB741 |
| TGCTAAACTTGCGAAAAATGTTTTTTTATTGCCGAGG TTAATCGATGAATTCGAGCTCG | This study | prMB742 |
| AGTGAACACTGAACAAGCATACTCTCAACCATTTAG ATACcggatccccgggttaattaa | This study | prMB830 |
| GTGACGATGATAACCGAGATGACGGAAATATAGTAC ATTAgaattcgagctcgtttaaac | This study | prMB831 |
| AGTGAACACTGAACAAGCATACTCTCAACCATTTAG ATACGGTCGACGGATCCCCGGG | This study | prMB832 |
| GTGACGATGATAACCGAGATGACGGAAATATAGTAC ATTAATCGATGAATTCGAGCTCG | This study | prMB833 |
| CGCTGAAGAAGAAGAAGTTG | This study | prMB1001 |
| GGAAGTCGCTGTTGCTAGTGCAAGCTCTTCCGCTTC AAGCcggatccccgggttaattaa | This study | prMB1002 |
| ACTTAAATGTTTCGCTTTTTTTAAAAGTCCTCAAAATT TAgaattcgagctcgtttaaac | This study | prMB1003 |
| GGAAGTCGCTGTTGCTAGTGCAAGCTCTTCCGCTTC AAGCcgtacgctgcaggtcgac | This study | prMB1004 |
| ACTTAAATGTTTCGCTTTTTTTAAAAGTCCTCAAAATT TAatcgatgaattcgagctcg | This study | prMB1005 |
| tctaagttttATGAAACACAATCGTCCAAATG | This study | prMB1363 |
| cacctttagaCATAATACTAAGCTGAAACAAAAAG | This study | prMB1364 |
| GAGGACTCAAAACGAAAAGG | This study | prMB1384 |
| GTGTTCTTTCTTCTGTACTATATATACATTTGCAACT ATGcggatccccgggttaattaa | This study | prMB1385 |
| TAAAAATTATAATGCCTAGTCCCGCTTTTGAAGAAAA TCAgaattcgagctcgtttaaac | This study | prMB1386 |
| TTACCAATTTAGCCACAAGG | This study | prMB1387 |
| ACCCTTGAAGATGGAGAATC | This study | prMB1388 |
| CGCCGAAACCCCCTCCCAAACTTCCCAAGAAGCAA CTCAAcgtacgctgcaggtcgac | This study | prMB1402 |
| TCGTGCGCTTTATTTGAATCTTATTGATCTAGTGAAT TTAatcgatgaattcgagctcg | This study | prMB1403 |
| ATTGTATAATATTACTCAACAGATTTTACAATTTTTAC ATcgtacgctgcaggtcgac | This study | prMB1406 |
| TAAAAATTATAATGCCTAGTCCCGCTTTTGAAGAAAA TCAatcgatgaattcgagctcg | This study | prMB1407 |
| CCGCTGAATAAGGTTTCTC | This study | prMB457 |
| AATAAAAAGTGACATCTGAAAAACATCCAATACTCC GATGcggatccccgggttaattaa | This study | prMB458 |

| | | |
|---|---|---|
| TGCAGATCTGATTTTTTTTCTATACTGTGCCTGTTCA TTAgaattcgagctcgtttaaac | This study | prMB459 |
| AGAAAAGAATGCGACGTATG | This study | prMB460 |
| GACAAAGAAGAAGCTGATGG | This study | prMB461 |
| CCAATGTGGGTGACTATGTGAG | This study | prMB481 |
| CTGTGGTTCCTGTAGTTGTG | This study | prMB482 |
| ACCACTTACAGGTGAGCAAC | This study | prMB801 |
| TAGGTCTAGCATCGTTTTCG | This study | prMB802 |
| acgctgtcagCATCGGAGTATTGGATGTTTTTCAG | This study | prMB545 |
| tactccgatgCTGACAGCGTTCAAAGTC | This study | prMB546 |
| cgaattcctgcagcccggggCATTACCCTAGACTTTCCTG | This study | prMB547 |
| ttgatagggaCATCGGAGTATTGGATGTTTTTCAG | This study | prMB548 |
| tactccgatgTCCCTATCAAAGAGTCATTTC | This study | prMB549 |
| cgaattcctgcagcccggggCATTACCCTAGACTTTCC | This study | prMB550 |
| aagaagaagcCATCGGAGTATTGGATGTTTTTCAG | This study | prMB551 |
| tactccgatgGCTTCTTCTTTGTCCGCC | This study | prMB552 |
| taggccccgaCATCGGAGTATTGGATGTTTTTCAG | This study | prMB554 |
| tactccgatgTCGGGGCCTATTACTGAG | This study | prMB555 |
| gccgaggttaGGCCATTTTGAAACTCATGTTG | This study | prMB576 |
| gccgaggttaTAGGGCCATTTTGTCGAGG | This study | prMB580 |
| cggccgctctagaactagtgGCGAAACTGAAGGTGGCAC | This study | prMB582 |
| gccgaggttaAACAGGTTCGAAAATAACATCTTCTTC | This study | prMB583 |
| cgaacctgttTAACCTCGGCAATAAAAAAAC | This study | prMB584 |
| acgctgtcagCGATGAATTCTCTGTCGG | This study | prMB592 |
| gaattcatcgCTGACAGCGTTCAAAGTC | This study | prMB593 |
| ttgatagggaCGATGAATTCTCTGTCGG | This study | prMB595 |
| gaattcatcgTCCCTATCAAAGAGTCATTTC | This study | prMB596 |
| aagaagaagcCGATGAATTCTCTGTCGG | This study | prMB598 |
| gaattcatcgGCTTCTTCTTTGTCCGCC | This study | prMB599 |
| cgaattcctgcagcccggggTTGTTGGTAGACTCAATGGC | This study | prMB560 |
| cggccgctctagaactagtgGAGCTCATAGCTTCAAAATG | This study | prMB648 |
| gccgaggttaGGCCATTTTGAAACTCATG | This study | prMB649 |
| caaaatggccTAACCTCGGCAATAAAAAAAC | This study | prMB650 |
| cgaattcctgcagcccggggTTGTTGGTAGACTCAATG | This study | prMB651 |
| gccgaggttaTAGGGCCATTTTGTCGAG | This study | prMB652 |
| aatggccctaTAACCTCGGCAATAAAAAAAC | This study | prMB653 |
| gccgaggttaAACAGGTTCGAAAATAACATC | This study | prMB654 |

### List of antibodies used in this study

| *Name* | *Source* | *Identifier* |
|---|---|---|
| Ldb16 | Bohnert lab | aMB11 |
| Ldo16/45 | Bohnert lab | aMB13 |
| Tdh1 | Abcam | Cat#125247 |
| Vac8 | Christian Ungermann | N/A |
| Pgk1 | Invitrogen | Cat#459250; RRID:AB_2532235g |
| GFP | Roche | Cat#11814460001 |
| anti-mouse IgG | Dianova | Cat#115-035-003 |

| | | |
|---|---|---|
| anti-rabbit IgG | Dianova | Cat#111-035-003 |
| anti FLAG M2 | Sigma | Cat#F3165; RRID:AB_259529 |
| anti-mouse IgG | Sigma | Cat#A4416; RRID:AB_258167 |
| anti-rabbit IgG | Sigma | Cat#A0545; RRID:AB_257896 |

#### List of software used in this study

| *Name* | *Source* | *Identifier* |
|---|---|---|
| Image J | Schneider et al., 2012 | https://imagej.nih.gov/ij/download.html |
| SerialEM | Mastronarde, 2005 | http://bio3d.colorado.edu/SerialEM/ RRID:SCR_017293 |
| IMOD | Kremer et al., 1996 | http://bio3d.colorado.edu/imod/ RRID:SCR_003297 |
| Topaz | Bepler et al., 2020 | https://github.com/tbepler/topaz |
| SNARTomo | This study | https://github.com/rubenlab/snartomo |
| MotionCor2 | Zheng et al., 2017 | https://emcore.ucsf.edu/ucsf-software RRID:SCR_016499 |
| Adobe Illustrator | Adobe Inc. | n/a |
| Affinity Photo | Serif | Version 1.7.3 RRID:SCR_016951 |
| Amira | Thermo Fisher Scientific | Version 2022.1 https://www.thermofisher.com/de/de/home/electron-microscopy/products/software-em-3d-vis/amira-software.html RRID:SCR_014305 |
| TomoSegMemTV | Martinez-Sanchez et al., 2014 | https://sites.google.com/site/3demimageprocessing/tomosegmemtv |
| Quick2D | Zimmermann et al., 2018; Gabler et al., 2020 | https://toolkit.tuebingen.mpg.de/tools/quick2d |
| Edge | Vilber Lourmat | Version 18.02 |
